# Cep55 overexpression promotes genomic instability and tumorigenesis in mice

**DOI:** 10.1101/780775

**Authors:** Debottam Sinha, Purba Nag, Devathri Nanayakkara, Pascal H.G. Duijf, Andrew Burgess, Prahlad Raninga, Veronique A.J. Smits, Amanda L. Bain, Goutham Subramanian, Meaghan Wall, John. W. Finnie, Murugan Kalimutho, Kum Kum Khanna

**Author notes:** These authors contributed equally (Murugan Kalimutho, Kum Kum Khanna). Correspondence (M.K), (K.K.K).

## Abstract

High expression of centrosomal protein CEP55 has been correlated with clinico-pathological parameters across multiple human cancers. Despite significant *in vitro* studies and association of aberrantly overexpressed CEP55 with worse prognosis, its causal role *in vivo* tumorigenesis remains elusive. Here, using a ubiquitously overexpressing transgenic mouse model, we show that *Cep55* overexpression causes spontaneous tumorigenesis and accelerates *Trp53^+/-^* induced tumours *in vivo*. At the cellular level, using mouse embryonic fibroblasts (MEFs), we demonstrate that *Cep55* overexpression induces proliferation advantage by modulating multiple cellular signalling networks including the PI3K/AKT pathway. Notably, the *Cep55* overexpressing MEFs demonstrate high level of mitotic chromosomal instability (CIN) due to stabilized microtubules. Interestingly, *Cep55* overexpressing MEFs have a compromised Chk1-dependent S-phase checkpoint, causing increased replication speed and DNA damage, resulting in a prolonged aberrant mitotic division. Importantly, this phenotype was rescued by pharmacological inhibition of Pi3k/Akt or expression of mutant Chk1 (S280A), that is insensitive to regulation by active AKT, in *Cep55* overexpressing cell. Collectively, our data demonstrates causative effects of deregulated Cep55 on genome stability and tumorigenesis which have potential implications for tumour initiation and therapy.

## Introduction

Genomic instability (GI) is a hallmark of almost all human cancers. Chromosomal instability (CIN) is a major form of GI, which refers to the acquisition of abnormal chromosome numbers or structures^1^. CIN in cancers primarily occur due to defective mitosis, including biased chromosome segregation and failure to undergo cytokinesis. Both mitotic checkpoint weakness and/or activation can also lead to CIN, exploring its genetic basis has the potential to uncover major mechanism of GI in cancers and therapeutic modality^2^.

CEP55 is a coiled-coil centrosomal protein which plays a critical role in cytokinetic abscission during mitotic exit^3^. CEP55 is a cancer testis antigen whose expression is restricted to male germ cells in adult animals, however it is re-expressed in a wide variety of cancers^4^. Over the last decade, multiple studies have shown variable associations of overexpressed CEP55 with poor prognosis in human cancers (reviewed by Jeffery *et al*. ^4^). On the other hand, loss-of-function mutations in *CEP55* cause Meckel-like and MARCH syndromes^5–8^. Notably, increased CEP55 expression correlates with functional aneuploidy in multiple cancer types, as defined by the *CIN70* gene signature^9^. It is also part of a 10-gene signature associated with drug resistance, CIN and cell proliferation^10^. Moreover, as part of the 31-gene cell-cycle progression (CCP) signature, it strongly correlates with actively proliferating prostate cancer cells^11^. Likewise, we have shown that *CEP55* is part of a 206 gene signature, representing genes enriched in promoting CIN, associated with aggressiveness in triple-negative breast cancer (TNBC)^12^.

Mechanistically, wild-type *TP53* suppresses CEP55 through PLK1 downregulation and therefore, cancers with *TP53* mutations often have elevated CEP55 levels^13^. In human cancers, CEP55 overexpression results in cell transformation, proliferation, epithelial-to-mesenchymal transition, invasion and cell migration via upregulation of the PI3K/AKT pathway through direct interaction with the p110 catalytic subunit of PI3K^14, 15^. Likewise, CEP55 interacts with JAK2 kinase and promotes its phosphorylation^16^. We have recently shown that *Cep55* overexpression causes male-specific sterility by suppressing Foxo1 nuclear retention through the PI3K/AKT pathway in mice^17^. Furthermore, we showed that CEP55 is a determinant of aneuploid cell fate during perturbed mitosis in breast cancers and could be targeted through MEK1/2-PLK1 inhibition^18^. Moreover, *Cep55* regulates spindle organisation and cell cycle progression in meiotic oocytes^19^. Collectively, these studies highlight the association of CEP55 overexpression with various human malignancies in a context-dependent manner. Though these *in vitro* and clinical correlation studies have so far established the link between CEP55 overexpression and cancer, the underlying mechanism by which CEP55 promotes tumorigenesis *in vivo* remains elusive.

Here, we report for the first time that *Cep55* overexpression in a mouse model causes high incidence of spontaneous tumorigenesis with a wide spectrum of highly proliferative and metastatic tumours. Notably, *Cep55* overexpression accelerates *Trp53^+/-^*-induced tumorigenesis. Using mouse embryonic fibroblasts (MEFs), we show that *Cep55* overexpression facilitates rapid proliferation by upregulating multiple cell signalling networks, particularly the PI3K/AKT pathway. Interestingly, we found that *Cep55* overexpression causes both numerical and structural CIN as a consequence of high frequency of anaphase chromatin bridges and micronuclei formation during chromosomal segregation with delayed mitotic exit due to stabilised microtubules. Mechanistically, *Cep55* overexpression compromised the Chk1-dependent S/G2 checkpoint due to hyperactivation of AKT signalling. As a consequence, the *Cep55* overexpressing MEFs cycle faster with increased replication fork speed, replication induced DNA damage and premature mitotic entry. Collectively, our data demonstrate a causal link of overexpressed Cep55 with tumorigenesis, driven through its multiple cellular functions.

## Results

### Cep55 overexpression drives tumorigenesis in vivo

To characterize the pathophysiological role of CEP55 overexpression *in vivo*, we utilised our recently reported transgenic mouse model^17^. Since *Cep55* is highly overexpressed in multiple human cancers irrespective of its role in cell division (Supp. Fig 1), we asked if *Cep55* overexpression causes spontaneous tumorigenesis *in vivo*. We monitored a cohort of wildtype (herein referred to as *Cep55^wt/wt^*, n=40), heterozygous transgenic (*Cep55^wt/Tg^,* n=40) and homozygous transgenic (*Cep55^Tg/Tg^*, n=50) *Cep55* mice over a period of 2.5 years for spontaneous tumour formation. We observed that the *Cep55^Tg/Tg^* mice developed various types of tumours at relatively long latencies (median survival 15 months) (Table 1) compared to other well-known oncogenic tumour models (K-ras^G12D 20^, Pten^+/- 21^ and Trp53^-/- 22, 23^). However, homozygous-*Cep55* overexpressing mice succumbed to cancer significantly earlier (p<0.0001) than *Cep55^wt/Tg^* and *Cep55^wt/wt^* littermates (Fig 1A). Notably, more than 50% of the *Cep55^Tg/Tg^* mice were culled in between 13-15 months due to tumour-associated phenotypes [irreversible weight loss (>15%), change in skin colour, reluctance to move and/or eat] (Supp. Fig 2A), suggesting that these mice might have developed tumours as a result of similar genetic changes caused by *Cep55* overexpression.

**Figure 1:**
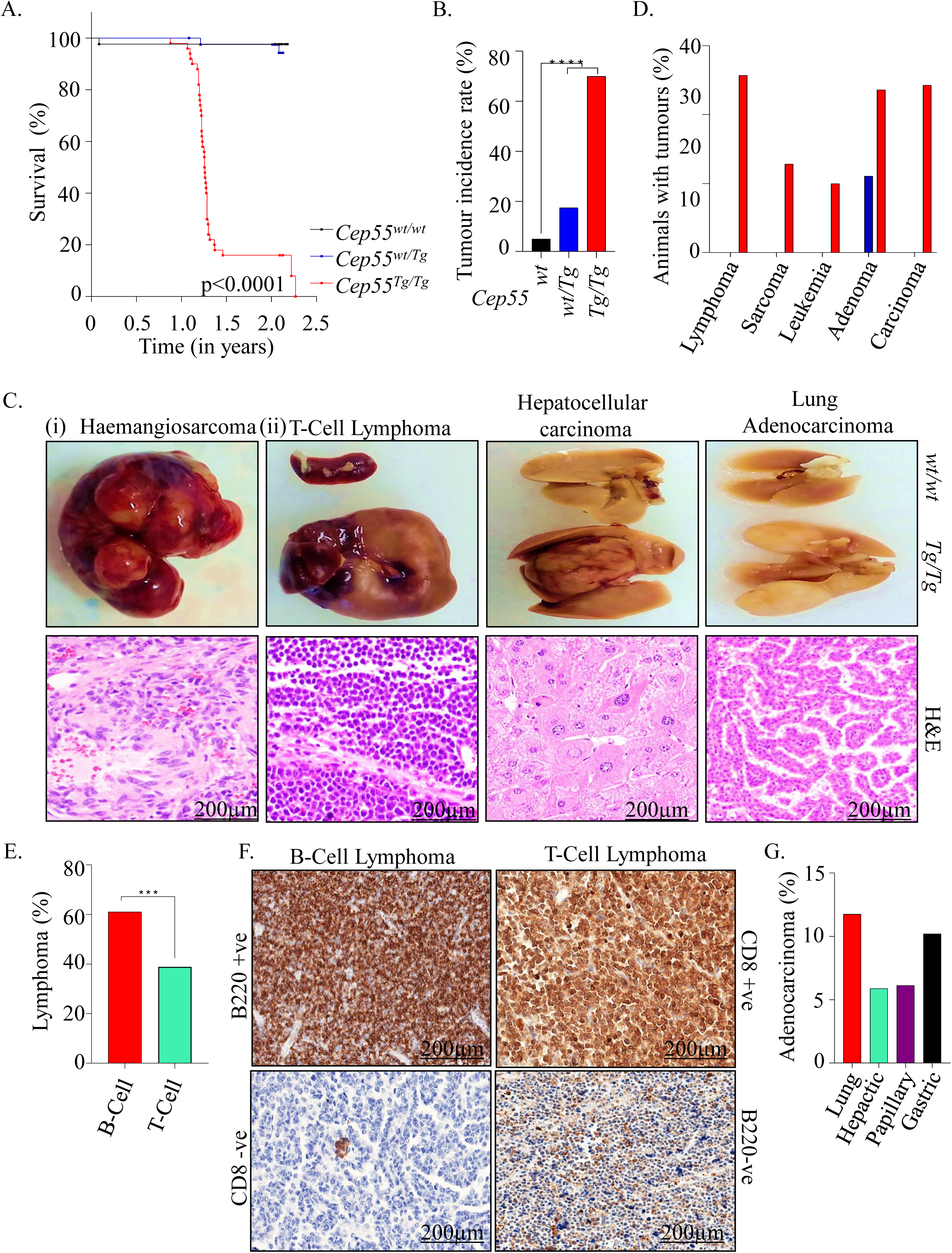
Cep55 overexpression causes spontaneous tumorigenesis in vivo. **(A)** Kaplan-Meier survival analysis of mice of indicated genotypes showing that *Cep55^Tg/Tg^* mice were more susceptible to form tumors compared to their control counterparts; Log-rank (Mantel-Cox) test was performed to determine *P-value* <0.0001. **(B)** Percentage of cancer incidence rate among mice of indicated genotypes (n≥40 per group); Fischer exact test was performed to determine *P-value*<0.0001 (****). **(C)** Representation images of gross morphology (upper panels) and H&E stained microscopic images (lower panels) of selected sections of (i) haemangiosarcoma in liver and (ii) indicated tumor lesions from different organs of tumor-bearing *Cep55^Tg/Tg^* mice (scale bars, 200µm). **(D)** Percentage of animals with respective cancer types observed in the transgenic cohorts. **(E)** Percentage of animal with types of lymphomas observed in the respective tumor bearing *Cep55^Tg/Tg^* mice. Fischer exact test was performed to determine P-value<0.0029 (***). **(F)** Representative images of B220 and CD8 immunostaining used to categorize the respective types of lymphomas. B220+ve and CD8-ve were classified as B-cell lymphoma while CD8+ve and B220-ve were classified as T-Cell lymphomas (scale bars, 200µm). **(G)** Percentage of adenocarcinoma in the respective organs observed in the tumors bearing *Cep55^Tg/Tg^* mice.

**Table 1:**
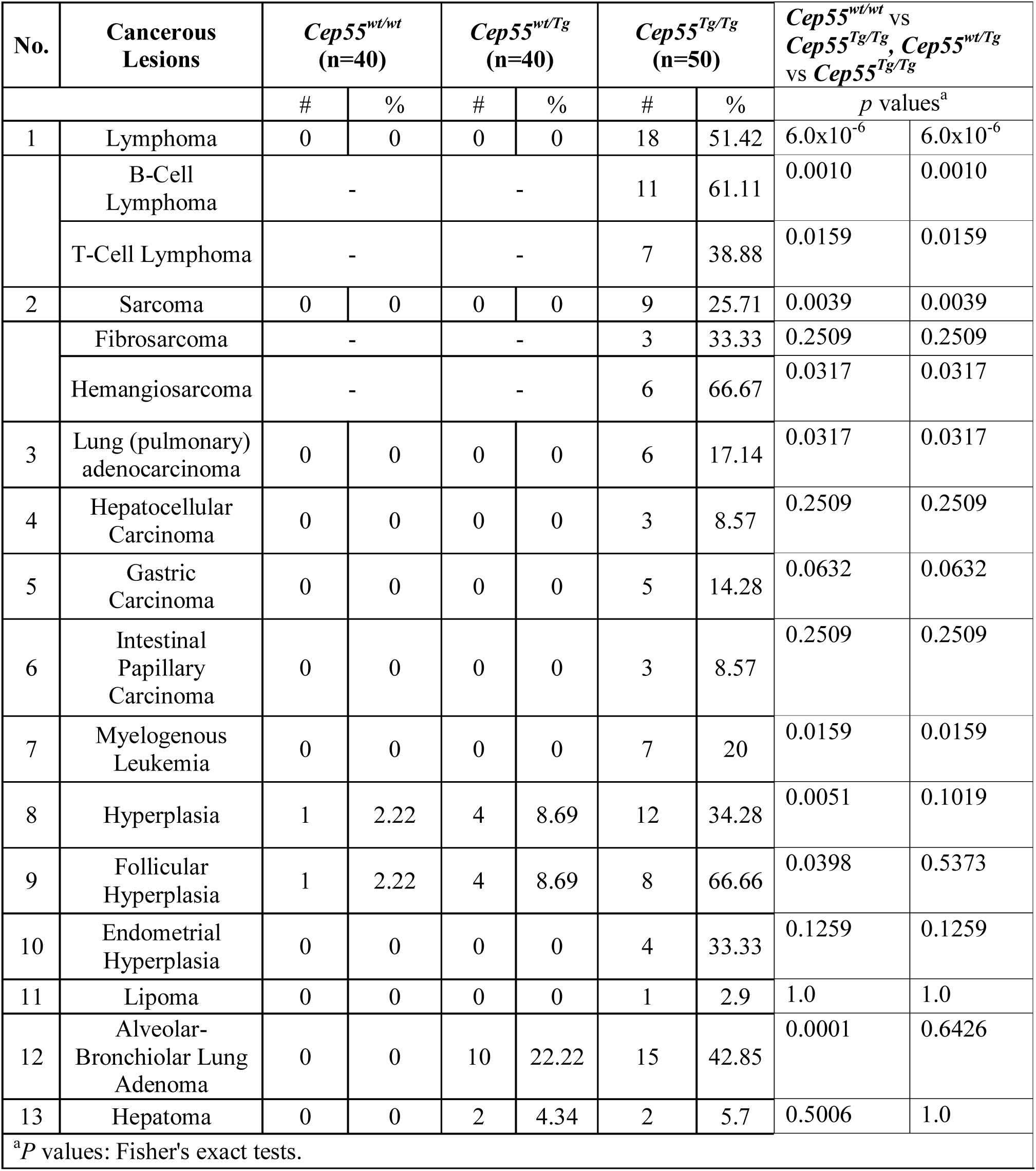
Distribution of cancer spectrum in *Cep55* transgenic mice.

We observed that 70% (35/50) of the *Cep55^Tg/Tg^* mice developed a wide spectrum of tumour lesions, including lymphoma, sarcoma, leukaemia and various adenocarcinomas (Fisher exact test p< 0.00001; Fig 1B-D, Supp. Fig 2B and Table 1) compared to only 17.5% (7/40) in *Cep55^wt/Tg^* and 5% (2/40) in *Cep55^wt/wt^* littermates (Fig 1B). Notably, the tumour burden observed in *Cep55^Tg/Tg^* mice varied between 1-3 tumours per animal (Supp. Fig 2C) with tumours originating in multiple tissue types (Supp. Fig 2D) in comparison to *Cep55^wt/Tg^*, which uniformly developed only adenomas in the lung. Likewise, the *Cep55^Tg/Tg^* mice also exhibited a higher incidence of lymphomas, in particular more B-cell lymphoma (1.5-fold) than the T-cell lymphoma (Fig 1D-E, Table 1). IHC staining using B220 (B-cell marker) and CD8 (T-cell marker) specified the incidence of B-cell and T-cell lymphoma’s, respectively (Fisher exact test p< 0.0029; Fig 1E-F). Independently, we observed a higher incidence of sarcomas, particularly more haemangiosarcoma than fibrosarcoma (in liver and spleen) (Supp. Fig 2D-E). We also observed a higher incidence of lung and gastric adenocarcinomas compared to other carcinomas (Fig 1G). We also observed a significant increase in hyperplastic lesions (in liver, spleen and endometrium) in *Cep55^Tg/Tg^* mice compared to the cohort of other genotypes (Fisher exact test p<0.0001; Supp. Fig 2F).

The primary cancers observed in the *Cep55^Tg/Tg^* mice were highly aggressive in nature with increased proliferation rate, as perceived by the gross morphology and mass of the organs in which these tumours originated (Fig 1C (ii), Supp. Fig 2G). In addition, we observed that ∼16% of the mice developed metastases (metastatic carcinoma) in the lungs and liver (Supp. Fig 2H) along with higher levels of inflammatory infiltrates, particularly lymphocytes (data not shown). Collectively, these data highlight that *Cep55* overexpression alone is sufficient to drive tumorigenesis in mice, causing a broad spectrum of cancers and associated abnormalities, such as inflammation and metastasis.

### Cep55 accelerates Trp53^+/-^ induced tumour development in mice

Our data suggests that *Cep55* overexpression-induced tumorigenesis mimics the tumorigenesis pattern observed in *Trp53^-/-^* mice^22^, as it induces a significantly higher percentage of lymphomas (∼35%) and sarcomas (∼17%) (Fig 1D). A previous report has demonstrated that wildtype TP53 restrains CEP55 expression through PLK1^13^. In addition, our clinical data mining suggests that *Cep55* levels are significantly higher in lung and hepatocellular tumours that exhibit allelic *TP53* copy number loss than in *TP53* diploid tumours (both p<0.0001, Mann-Whitney *U* test) (Supp. Fig 3A). Consistent with this, we observed a high p53 protein level, which is most likely an indication of mutated *Trp53*, in representative *Cep55^Tg/Tg^* tumour tissues than normal adjacent tissues (Fig 2A).

**Figure 2:**
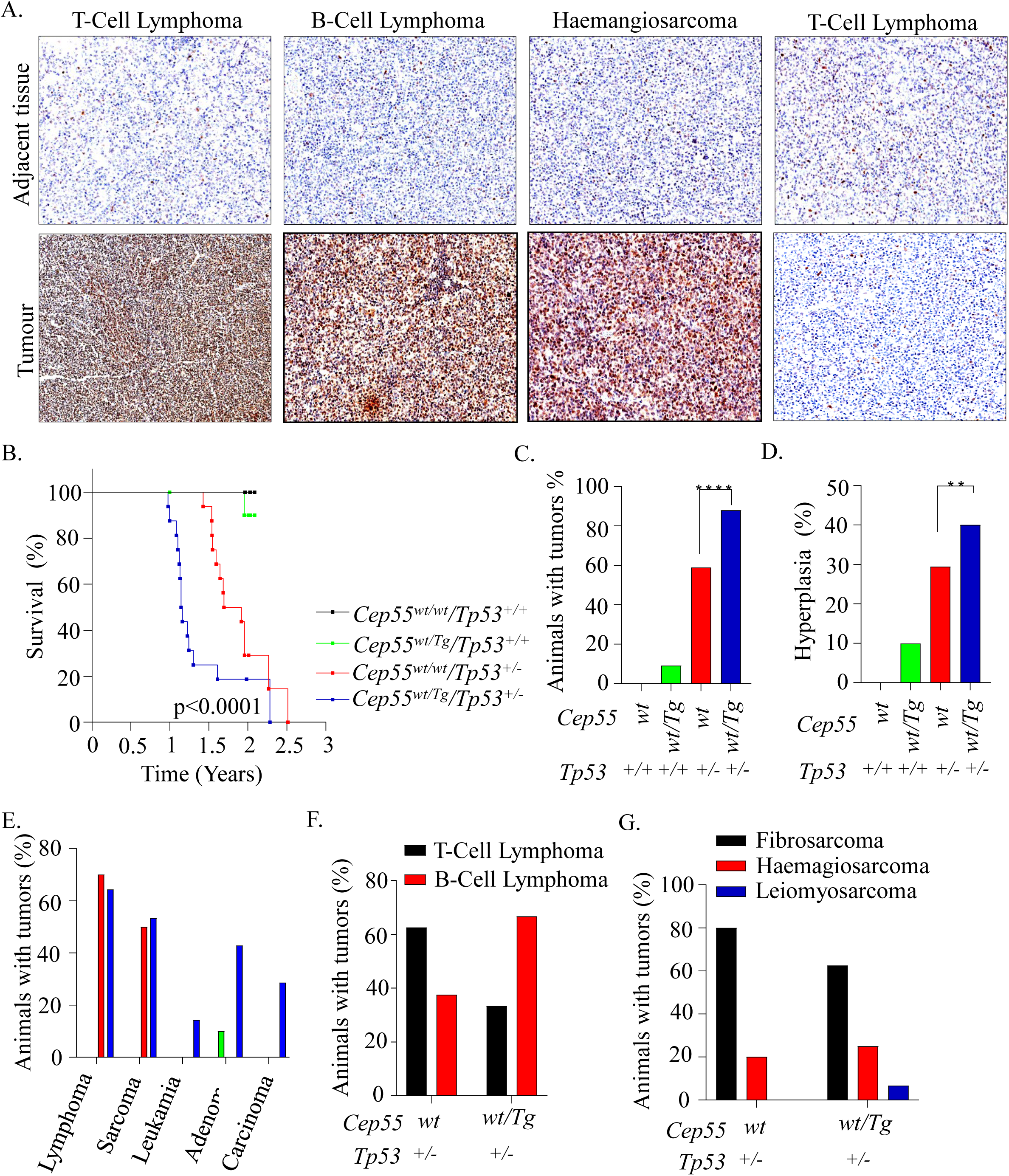
Heterozygous Cep55 transgenic expression accelerates Trp53^+/-^ induced tumorigenesis in mice. **(A)** Representative images of p53 immunohistochemical staining on tumor sections of respective subtypes observed in the *Cep55^Tg/Tg^* mice (bottom panel) in comparison to adjacent normal tissue from the same mice (upper panel). **(B)** Kaplan-Meier survival analysis highlighting the tumor-free survival of the mice of indicated genotypes demonstrating that the *Cep55^wt/Tg^; Trp53^wt/-^* mice were more susceptible to form tumors with a shorter latency period (∼14 months) compared to control counterparts; Log-rank (Mantel-Cox) test was performed to determine *P-value* <0.0001. **(C-G)** Percentages of overall cancer incidence **(C),** hyperplastic lesions **(D),** cancer spectrum **(E),** lymphoma **(F)** and sarcoma burden **(G)** among mice of indicated genotypes (n≥ group). Fischer exact test was performed to calculate *P-value* <0.0001 (****) and <0.01(**).

Next, to examine the contribution of *Cep55* overexpression to *Trp53^+/-^*-induced tumorigenesis, we inter-crossed *Cep55^Tg/Tg^* female mice with *Trp53^-/-^* male mice to establish bi-transgenic cohorts of *Cep55^wt/Tg^;Trp53^+/-^* (n=15), *Cep55^wt/wt^;Trp53^+/-^* (n=17), *Cep55^wt/Tg^;Trp53^+/+^* (n=11) and *Cep55^wt/wt^;Trp53^+/+^* (n=10) mice. These cohorts of mice were monitored regularly for a period of 2.5 years for spontaneous tumour development. Interestingly, we observed that the *Cep55^wt/Tg^;Trp53^+/-^* mice succumbed to a broad spectrum of cancer development (spleen, liver and lung) with reduced latency (median survival of 13.8 months; p<0.0001) when compared to the *Cep55^wt/wt^;Trp53^+/-^* cohort (median survival of 21.6 months) (Fig 2B, Supp. Fig 3B-F, Supp. Table 1). Notably, the entire cohort of *Cep55^wt/Tg^;Trp53^+/-^* mice exhibited a time frame of tumour development similar to that of *Cep55^Tg/Tg^* mice (Fig. 2B), suggesting a strong contribution of Cep55 in inducing *Trp53^+/-^* mediated tumours.

Further, the incidence of tumorigenesis observed in *Cep55^wt/Tg^;Trp53^+/-^* mice was also significantly higher (∼85%; Fisher exact test p<0.0001) in comparison to *Cep55^wt/wt^;Trp53^wt/-^* (∼50%) with 1-3 tumours per animal (Fig 2C). The *Cep55^wt/Tg^;Trp53^+/-^* mice also displayed a significantly higher incidence of hyperplastic lesions (Fisher exact test p<0.01) (Fig 2D), and a similar incidence to that observed in *Cep55^Tg/Tg^* mice (Supp. Fig 2F). Histopathological analysis indicated the presence of a number of neoplastic lesions (Fig 2E, Supp. Fig 3B, Supp. Table 1) that were similarly observed in *Cep55^Tg/Tg^* mice (Fig 1C-D, Supp. Fig 2A and Table 1). Notably, though a similar fractions of *Cep55^wt/wt^;Trp53^+/-^* and *Cep55^wt/Tg^;Trp53^+/-^* animals developed lymphomas and sarcomas (Fig. 2E), their lymphoma spectrums were different, in particular there was a higher incidence of B-cell lymphomas than T-cell lymphomas in the *Cep55^wt/Tg^;Trp53^+/-^* mice compared to *Cep55^wt/wt;^Trp53^+/-^* mice (Fig 2F). Further, the *Cep55^wt/Tg^;Trp53^+/-^* mice demonstrate a similar occurrence of fibrosarcoma and haemangiosarcoma (in liver and spleen), as observed in *Cep55^Tg/Tg^* mice (Fig 2G). Taken together, these data indicate that *Cep55* overexpression accelerates tumourigenesis and changes the tumour spectrum in *Trp53^+/-^* mice.

### Cep55 overexpression confers a survival advantage through activation of signalling networks

In multiple human cancers, deregulated expression of *CEP55* has been linked to enhanced proliferation, migration, invasion, epithelial-mesenchymal transition and tumorigenesis^4^. To analyse the cellular consequences of *Cep55* overexpression *in vivo*, we used primary and spontaneously immortalized MEFs generated from our transgenic mice (Supp. Fig 4A). We observed significantly higher *Cep55* transcript and protein levels in the *Cep55^Tg/Tg^* MEFs compared to MEFs from other genotypes (Fig 3A-B). Next, to determine the growth potential and the senescence rate in the primary MEFs, we performed an NIH-3T3 assay and observed that the *Cep55^Tg/Tg^* primary MEFs had a significantly higher proliferation rate in comparison to *Cep55^wt/Tg^* and *Cep55^wt/wt^* MEFs, with an increased G2/M proportion of cells (Fig 3C-D). However, no statistically significant difference was observed in the proliferation rates between *Cep55^wt/Tg^* and *Cep55^wt/wt^* (Fig 3C). Likewise, the immortalized *Cep55^Tg/Tg^* MEFs also exhibited similar enhanced proliferative capacity with a significant difference in cell cycle distributions (Fig 3E-F, Supp. Fig 4B). To define if *Cep55* overexpression alone could confer enhanced proliferative capacity independent of mitogenic signals, we serum-starved the primary MEFs of each genotypes and observed higher cell proliferation capacity in *Cep55^Tg/Tg^* MEFs (∼60 hrs) compared to MEFs from other genotypes, highlighting a self-mitogen gaining capability to proliferate and survive in conditions of serum-starvation (Supp. Fig 4C).

**Figure 3:**
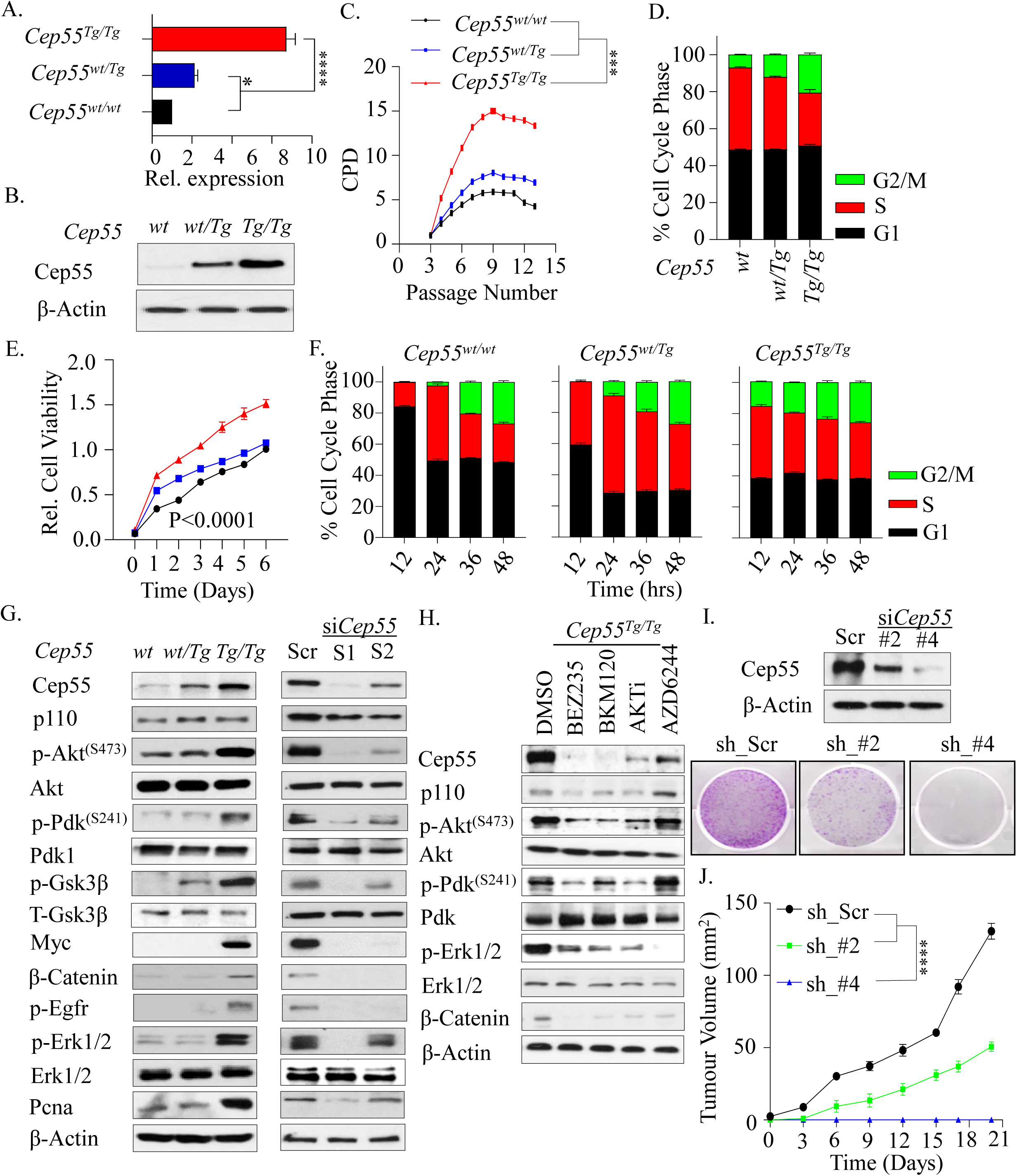
Cep55 confers survival advantage through signaling networks. **(A)** Statistical representation of transgenic expression of *Cep55* transcripts observed in the mouse embryonic fibroblasts (MEFs) of respective genotypes (n=3 per group). One-way ANOVA test was performed to determine *P-value* <0.0001 (****) and <0.05(*). **(B)** Immunoblot analysis of Cep55 expression in the whole cell lysates of the primary MEFs of each genotype. β-Actin was used as loading control. **(C)** Statistical representation of NIH-3T3 proliferation assay measured as a function of passage number [indicated as CPD (cumulative population density)] observed in primary *Cep55^Tg/Tg^* MEFs in comparison to its counterparts (n=3 per group). One-way ANOVA test was performed to determine *P-value* <0.0001 (****). **(D)** Statistical representation of the cell cycle profile of primary MEFs of indicated genotype at 24 hours (n=3 per group). **(E)** Statistical representation of the cell viability of immortalized MEFs of each genotype, as indicated in (C), measured per day over a period of 6 days (n=3 per group). One-way ANOVA test was performed to determine *P-value* <0.0001 (****). **(F)** Comparison of cell cycle profile of immortalized MEFs of the indicated genotype over 48 hours measured at 12-hour intervals (n=3 per group). **(G)** Immunoblot analysis of the whole cell lysates collected after 24 hours from the immortalized MEF’s of indicated genotypes (left panel) and 48 hours from the respective siRNA treated *Cep55^Tg/Tg^* MEFs (right panel) indicating the impact of *Cep55* overexpression on multiple cell signaling pathways. Β **(H)** Immunoblot analysis of the whole cell lysates collected after 24 hours of treatment with the respecting inhibitors. β-Actin was used as loading control **(I)** Immunoblot analysis of the whole cell lysates collected from the respective isogenic *Cep55*-depleted TCLs at 24 hours validating the levels of Cep55 expression. β-Actin was used as loading control (left panel). Representative images of colony formation at 14 days determined using crystal violet staining in control and *Cep55*-depleted TCLs (right panel). **(J)** Six-week-old female NOD/Scid cohorts of mice were injected subcutaneously with the control and *Cep55*-depleted clones. Growth rates (area, mm^2^) of the tumors were measured using a digital caliper. Differences in growth were determined using Student’s t-test, P ≤ 0.0001 **(****)**. Graph represents the mean tumor area ± SEM, n = 5 mice/group.

CEP55 has been shown to upregulate AKT phosphorylation through direct interaction with p110 catalytic subunit of PI3 kinase (PI3K) and enhance cell proliferation *in vitro*^14, 15, 17^. Likewise, we have shown that MYC regulates CEP55 transcriptionally in breast cancer^18^. Thus, to characterize the molecular signalling involved in cell proliferation and survival, we investigated the impact of *Cep55* overexpression on Pi3k/Akt - and Erk-dependent signalling networks. Interestingly, immunoblot analysis using whole cell lysates from the MEFs of each genotype demonstrated *Cep55* allele-dependent increase in phosphorylation of Akt^S473^ and its upstream regulator Pdk1^S241^ in *Cep55^Tg/Tg^* MEFs compared to wild type and heterozygous MEFs (Fig 3G). In addition, we also observed an upregulation of Mapk-dependent signalling molecules, including increased-phosphorylation of Egfr, Erk1/2, Myc and β-catenin, along with increased Pcna, a proliferation marker, in *Cep55^Tg/Tg^* MEFs (Fig 3G). Similar changes were observed in representative tissue lysates (Supp. Fig 4D). Notably, the effects on the signalling networks were specific to *Cep55* overexpression as knockdown of *Cep55* using two different siRNA oligonucleotides in *Cep55^Tg/Tg^* MEFs remarkably diminished Pi3k/Akt and Mapk-dependent signalling pathway activities (Fig 3G) and proliferation rate (data not shown). Furthermore, to characterize the role of *Cep55* overexpression in promoting cell proliferation and survival through activated signalling pathways, we used a wide range of Pi3k/Akt, mTor and Erk1/2 pathway-specific inhibitors. Blocking these signalling pathways markedly reduced Cep55 levels suggesting a positive feedback loops between Cep55 and these signalling pathways (Fig 3H). Moreover, we observed that the *Cep55^Tg/Tg^* MEFs were significantly more sensitive to AKT, PI3K and pan-PI3K-AKT/mTOR inhibitors, but not to mTOR or Erk1/2 inhibitor treatments alone (Supp. Fig 4E), suggesting a higher dependency of these cells on Pi3k/Akt signalling.

To further decipher the impact of overexpressed *Cep55* on tumorigenesis, we established cell lines from some of the tumours that developed in *Cep55* overexpressing mice (herein abbreviated as tumour cell lines (TCLs)), in particular haemangiosarcoma of the liver (Fig 1C(i)). These cells exhibited a mixed population of bi- and multi-nucleated cells, implying a genomically unstable phenotype (Supp. Fig 4F). Notably, upon transient *Cep55* knockdown using siRNA in the TCL, these cells tended to grow slower with a concomitant reduction in Pdk1 and Akt phosphorylation levels. We only observed a marginal Myc reduction, while there was no impact on Erk1/2 levels (Supp. Fig 4G). Likewise, constitutive *Cep55* knockdown in this line using shRNAs reduced anchorage-independent colony formation (Fig 3I, Supp. Fig 4H), proliferation rate and tumour formation dependent on the extent of reduction of Cep55 levels (Fig 3J, Supp. Fig 4I). Consistently, the *Cep55* knockdown TCL were significantly refractory to PI3K/Akt inhibitor sensitivity (Supp. Fig 4K), suggesting a dependency on PI3K/Akt signalling. Taken together, these data highlight the crucial role of *Cep55* in regulating proliferation and survival-associated signalling networks and its essential function in tumour formation.

### Cep55 overexpression promotes structural and numerical chromosomal instability (CIN)

The well-known role of CEP55 as a regulator of CIN is through regulation of cytokinesis^3^. Consistent with this, we found that whole-genome duplicated (WGD) tumours have significantly higher levels of *CEP55* mRNA than diploid and near diploid tumours (Supp. Fig 5A). Likewise, *Cep55^Tg/Tg^* MEFs exhibited a three-fold higher percentage (p<0.0001) of binucleated and multinucleated cells (Fig 4A-B). In addition, using FACS analysis, we found that both primary and immortalized *Cep55^Tg/Tg^* MEFs exhibited a significantly higher percentage of >4n subpopulation (Fig 4C, Supp. Fig 5B). Similar results were observed in different organs isolated from *Cep55^Tg/Tg^* mice compared to their littermate counterparts (Supp. Fig 5D). Importantly, we found a significant increase in micronuclei in the *Cep55^Tg/Tg^* MEFs (p<0.001) indicating the possible presence of CIN (Fig 4D). Likewise, when *Cep55* was constitutively knocked down in TCLs, we found a significant reduction in >4n subpopulations (Fig 4E-F, Supp. Fig 5E), suggesting that Cep55 overexpression facilitates CIN. Consistent with this, when we analysed the level of aneuploidy across some of the human cancers using using Genome-wide SNP6 array data from TCGA, we found that *CEP55* overexpressing tumours show increased structural or numerical aneuploidy, including whole-chromosome aneuploidy and chromosome arm-level aneuploidy (Supp. Fig 6A-D). Additionally, spectral karyotyping of metaphase spreads from *Cep55^Tg/Tg^* MEFs demonstrated the presence of significantly higher levels of both numerical and structural chromosomal aberrations compared to other genotypes (Fig 4G). Notably, these MEFs demonstrated complex chromosomal translocations and numerical abnormalities, wherein both *Cep55^wt/Tg^* and *Cep55^wt/wt^* MEFs showed a low level of structural and numerical chromosomal abnormalities (Table 2). In summary, these data highlight that Cep55 overexpression above a certain threshold is sufficient to promote structural and numerical CIN *in vitro* as well as *in vivo*.

**Figure 4:**
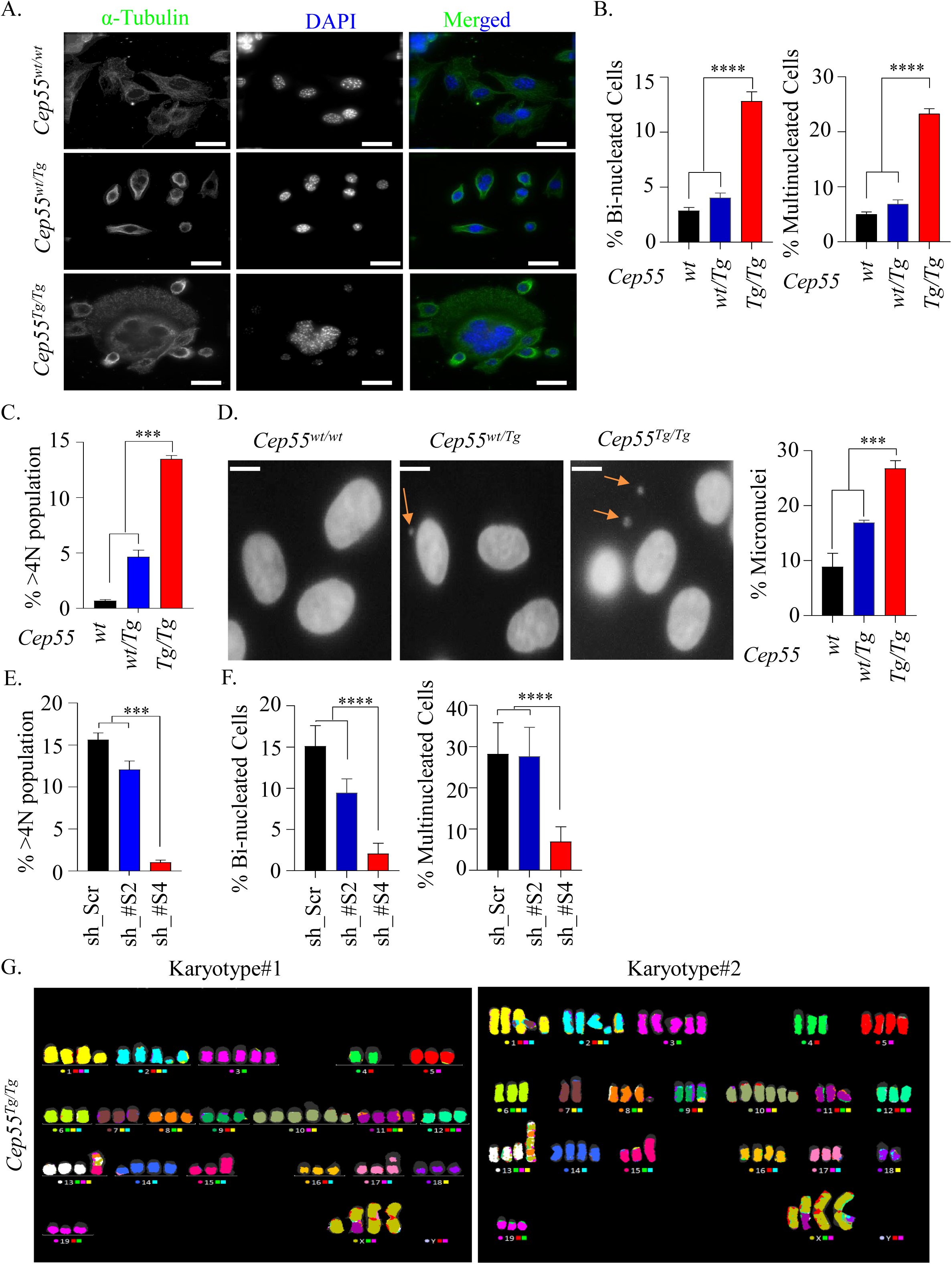
Cep55 overexpression promotes chromosomal instability in vivo. **(A)** Representative images of immunofluorescence demonstrating genomic instability observed in *Cep55^Tg/Tg^* MEFs, as indicated by the presence of multiple nuclei (marked by DAPI staining) compared to other counterparts. The cell cytoplasm is marked by α-tubulin (green) (Scale bar, 100 μm). **(B)** Statistical representation showing the percentage of binucleated (left panel) and multinucleated cells (right panel) observed in the respective immortalized MEFs (n=100 cells of each genotype). Error bars represent the ± SD from two independent experiments. One-way ANOVA test was performed to determine *P-value* <0.0001 (****). **(C)** Statistical representation of polyploidy analysis (>4N DNA contents) determined using FACS analysis in the respective immortalized MEFs. Error bars represent the ± SD from two independent experiments. One-way ANOVA test was performed to determine *P-value* <0.001 (***). **(D)** Representative images showing the presence of micronuclei (marked by DAPI) in the respective immortalized MEFs (left panel) (Scale bar, 100 μm). Statistical representation showing the percentage of micronuclei observed in the respective immortalized MEFs (right panel). Error bars represent the ± SD from two independent experiments. One-way ANOVA test was performed to determine *P-value* <0.001 (***). **(E)** Statistical representation of polyploidy analysis (>4N DNA contents) determined using FACS in the respective sh*Cep55* depleted isogenic clones. Error bars represent the ± SD from two independent experiments. One-way ANOVA test was performed to determine *P-value* <0.001 (***). **(F)** Statistical representation showing the percentage of binucleated (left panel) and multinucleated cells (right panel) observed in the respective sh*Cep55* depleted isogenic clones. (n=100 cells per clone). Error bars represent the ± SD from two independent experiments. One-way ANOVA test was performed to determine *P-value* <0.0001 (****). **(G)** Representative metaphases from spectral karyotyping (SKY) in the *Cep55^Tg/Tg^* MEFs (passage 25) wherein #1 and #2 denotes biologically independent metaphase representation of *Cep55^Tg/Tg^* MEFs.

**Table 2:** Changes in chromosomal alterations in *Cep55* transgenic MEFs.

| Genotype | Karyotype | Phenotype |
| --- | --- | --- |
| *Cep55<sup>wt/wt</sup>* | 77,XXXX,-6,-7,-18[17] | Hypotetraploid with numerical abnormalities. |
| *Cep55<sup>wt/Tg</sup>* | 80,XXXX[6]/77,idem,-6,-7,-18[11]/40,XX[4] | Four normal female metaphases. Six tetraploid metaphases and eleven hypotetraploid metaphases with the same numerical abnormalities that were seen in the WT cell line. |
| *Cep55<sup>Tg/Tg</sup>* | 72~74,X,der(X)t(X;11)(F?1;A?2),i(X)(A1)x2,del(1)(A?E?),del(2)(?B?H),+3,-4,-6,-7,del(8)(A?2),-9,der(9)(9pter->9?F::2??2?F::1?H>1qter)[3],der(9)t(9;17)(F?;E?1)[2],+10,+10,del(10)(A2B4)x3,-11,-12,der(13)(13pter->13?::8?->8?::13?->13?:: 8?->8?:: 13?->13qter)[12],der(13) (13pter->13?::8?->8?::13?->13?:: 8?->8?:: 13?->13?::5?->5qter)[2],<br>der(13) (13pter->13?::8?->8?::13?->13?:: 8?->8?:: 13?->13?::15?-><br>5qter)[3],der(13)t(13;14)(A?;B?)[2],-15,dup(15)(ED?2),-17,der(17)<br>t(9;17)(?F1;?B)[3],i(17)(A1),-18,-19[cp17] | Hypotetraploid with complex numerical and structural abnormalities. |

Note: ? = questionable identification of chromosome or chromosome structure

### Cep55 overexpression delays mitotic exit

CIN in cancers primarily occurs due to defective mitosis including unequal chromosome segregation and failure to undergo cytokinesis. Our initial analysis of percentage of cells undergoing mitosis revealed that *Cep55^Tg/Tg^* MEFs had a significantly increased mitotic index compared to other genotypes (Supp. Fig 7A-B; p<0.001) and *Cep55*-depleted TCLs showed a reduction in the number of mitotic cells (Supp. Fig 7C). We next asked how *Cep55* overexpression might promote both structural and numerical CIN in these cells during normal and perturbed mitosis. To decipher this, we collected double-thymidine synchronised MEFs for DNA content and time-lapse live-cell imaging analyses. Notably, we observed that the *Cep55^Tg/Tg^* MEFs progressed faster through interphase and entered mitosis more rapidly compared to *Cep55^wt/wt^* MEFs (Supp. Fig 7D). However, the *Cep55^Tg/Tg^* MEFs spent a relatively longer time in mitosis with a higher percentage of cells exhibiting cytokinesis failure compared to wildtype and heterozygous MEFs (Fig 5A-B). Likewise, the *Cep55^wt/Tg^* MEFs also spent significantly more time in mitosis compared to wildtype MEFs, indicating an allele-dependent impact of *Cep55* overexpression on mitotic duration (Fig 5A). Multinucleated cells usually take more time to complete mitosis due to high DNA content and the *Cep55^wt/Tg^* and *Cep55^Tg/Tg^* MEFs exhibited mixed subpopulations of mononucleated, binucleated and multinucleated cells (Fig 4B, C). We therefore performed analysis of individual subpopulations to determine the duration of mitosis (Fig 5C). Surprisingly, along with the bi- and multinucleated *Cep55^Tg/Tg^* cells, the mononucleated cells also spent more time in mitosis, indicating that *Cep55* overexpression prolonged mitotic duration independently of DNA content (Fig 5D). Moreover, consistent with our previous report in breast cancer^18^, *Cep55* overexpression significantly impacted the duration of time to- and time spent in mitosis upon nocodazole treatment (Fig 5E-F). In particular, the *Cep55^Tg/Tg^* cells largely prematurely exited mitosis during nocodazole arrest but the *Cep55^wt/wt^* cells predominately died in mitosis (Fig 5G-J). On contrary, the *Cep55*-depleted TCL showed sensitivity towards nocodazole treatment with significant reduction in premature exit and increase in apoptosis (Supp. Fig 7F-H). Therefore, these data indicate that *Cep55* overexpression facilitates CIN through interference with normal cell cycle regulation.

**Figure 5:**
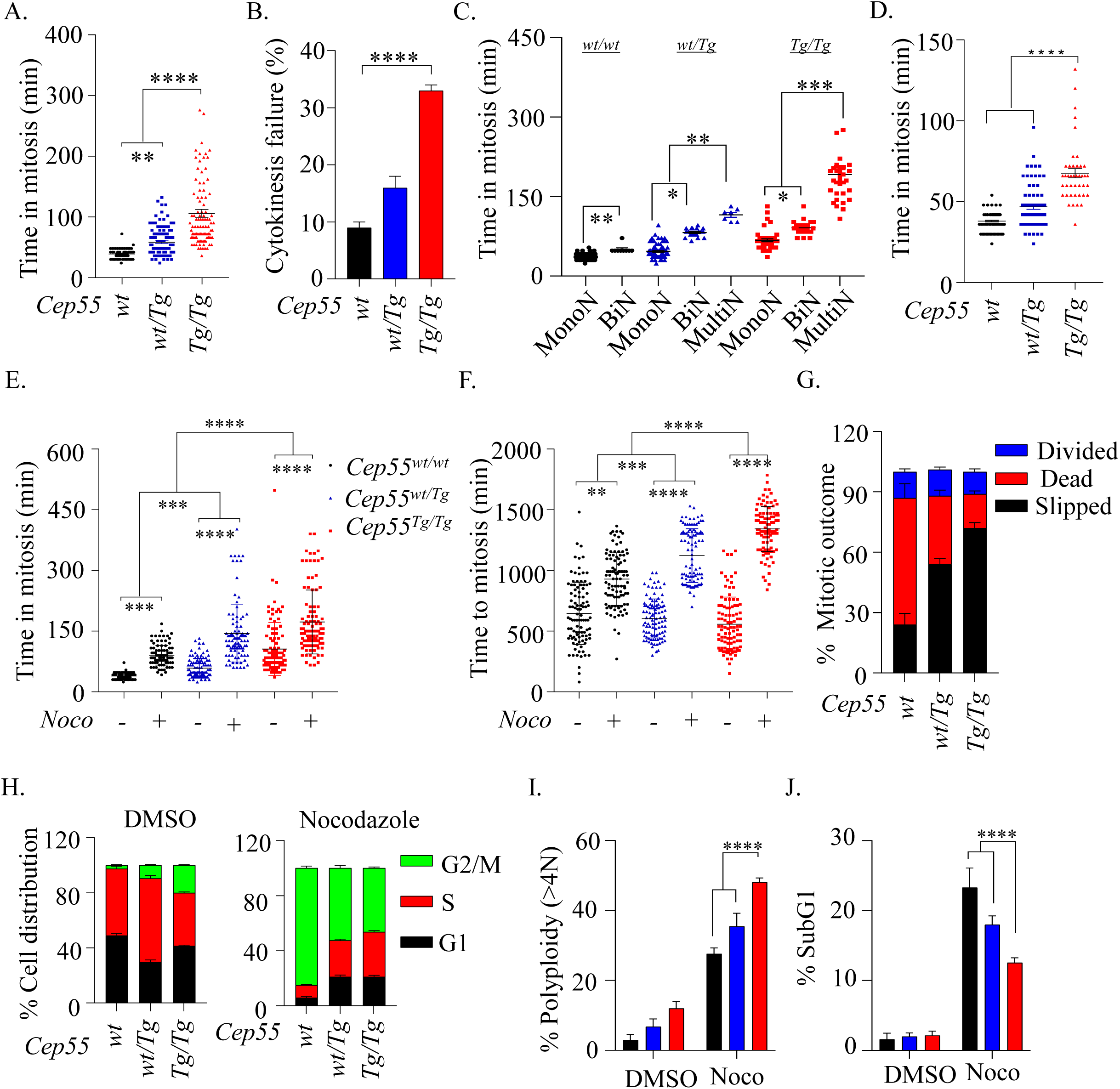
Impact of Cep55 overexpression on mitosis. **(A)** Statistical representation showing average time spent in mitosis by the MEFs of indicated genotypes. The MEFs were synchronized using double-thymidine block and released in regular culture media. Individual cells were tracked using bright-field Olympus Xcellence IX81 time-lapse microscopy for overall time taken to complete mitosis from nuclear envelope breakdown up to daughter cell formation, as described previously^18^. **(B)** Statistical representation showing the percentage of cytokinesis failure observed in the MEFs of indicated genotypes. **(C)** Statistical representation showing average time spent in mitosis by the different cell population observed among the MEFs of indicated genotypes. **(D)** Statistical representation showing average time spent in mitosis by mononucleated MEFs of indicated genotypes. **(E)** Statistical representation showing comparison of average time spent in mitosis by MEFs of indicated genotypes in presence or absence of nocodazole (0.5 μM) Time in mitosis was calculated as in (A). **(F)** Statistical representation showing comparison of average time taken by the MEFs of indicated genotypes to enter mitosis after release from double-thymidine block, presence and absence of nocodazole (0.5 μM). Time to enter mitosis was calculated as in (A). Error bars represent the ± SD from two independent experiments of all the above expreiments. One-way ANOVA test was performed to determine *P-value* <0.05 (*), <0.01 (**), <0.001 (***) and <0.0001 (****). **(G)** Statistical representation showing the mitotic outcome in the MEFs of indicated genotypes in presence of nocodazole (0.5 μM). Mitotic slippage was defined by premature mitotic exit during nocodazole-induced mitotic arrest, while death was determined through membrane blebbing. **(H)** Statistical representation of the cell cycle profiles of MEFs of indicated genotype in the presence or absence of *nocodazole (0.5* μ*M)* (n=2 per group). **(I)** Statistical representation of polyploidy analysis (>4N DNA contents) determined using FACS in the indicated immortalized MEFs in the presence or absence of *nocodazole (0.5* μ*M)*. Error bars represent the ± SD from two independent experiments. One-way ANOVA test was performed to determine *P-value* <0.0001 (****). **(J)** Statistical representation of percentage SubG1 populations was determined using FACS in the indicated immortalized MEFs in the presence or absence of *nocodazole (0.5* μ*M)* (n=3 per group). Error bars represent the ± SD from two independent experiments. One-way ANOVA test was performed to determine *P-value* <0.0001 (****).

### Cep55 overexpression induces defective chromosomal segregation due to stabilised microtubules

Chromosome segregation errors are a major source for CIN^24^. As we observed that *Cep55^Tg/Tg^* MEFs demonstrate a higher rate of CIN and homozygous *Cep55-*overexpressing cells spend more time in mitosis compared to cells of other genotypes (Fig 5A-D), we next investigated the impact of *Cep55* overexpression on chromosome segregation during mitosis. Double-thymidine synchronized cells were fixed after ∼6hrs post-release and mitotic cells were visualised using fluorescence microscopy. Surprisingly, *Cep55^Tg/Tg^* MEFs demonstrated a significantly higher frequency (p<0.05) of multipolar spindle poles along with unaligned and lagging chromosomes compared to *Cep55^wt/wt^* MEFs (Fig 6A-D, Supp. Fig 8A-C). In addition, using both fluorescence and live-cell time-lapse microscopy, we also observed that the *Cep55^Tg/Tg^* MEFs showed significantly higher frequency of anaphase cells with chromatin bridges (anaphase bridges). The presence of anaphase bridges during mitosis indicates the presence of incompletely segregated DNA in *Cep55^Tg/Tg^* MEFs which in turn result in chromosomal breakage and micronuclei formation (Fig 6E, Supp. Fig 8D). Consistent with this, we observed that the *Cep55^Tg/Tg^* MEFs exhibited an increased proportion (p<0.001) of micronuclei, a morphological characteristic of CIN, when compared to control MEFs (Fig 6F, Supp. Fig 8D).

**Figure 6:**
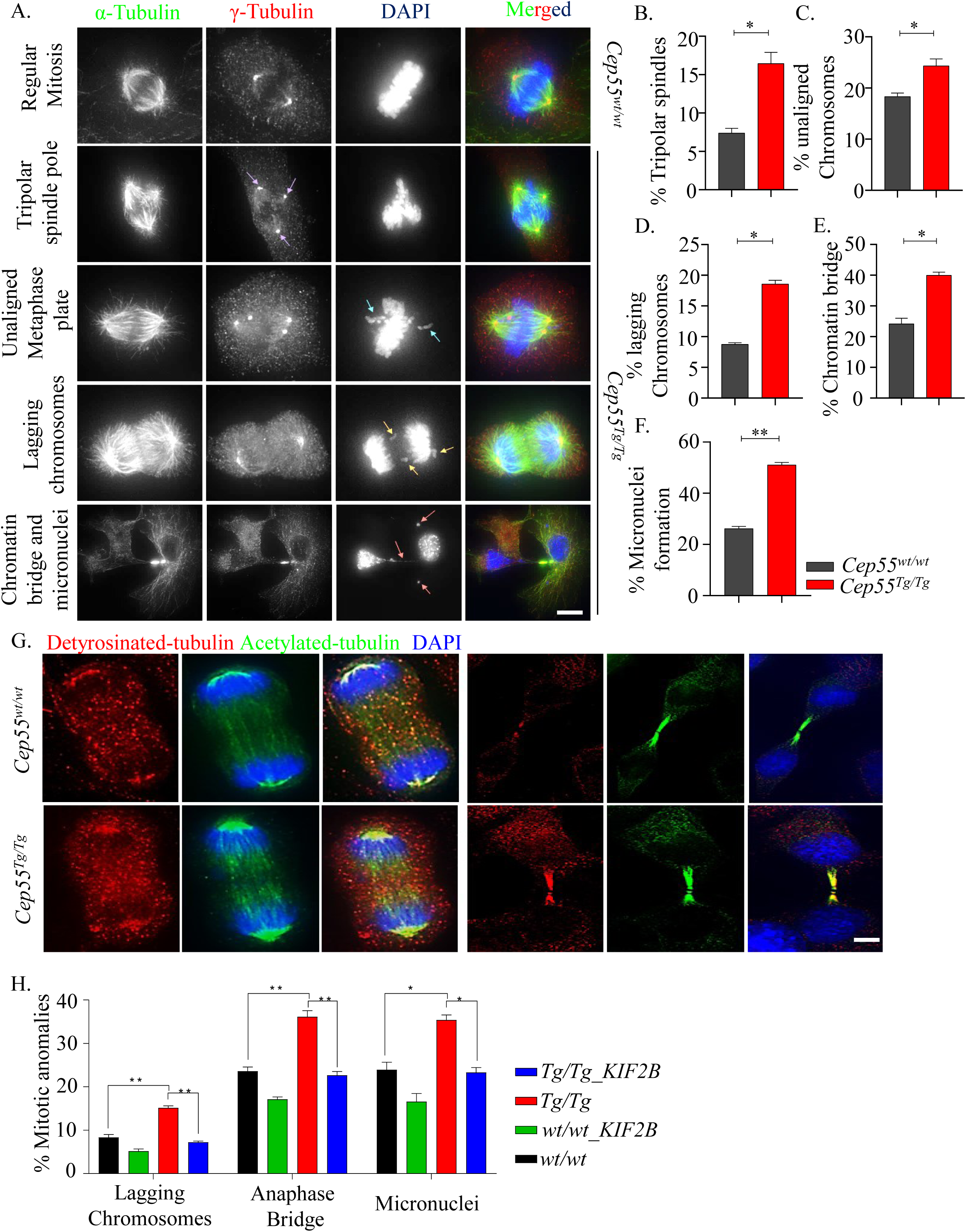
Cep55 overexpression causes various mitotic defects. **(A-F)** Representative images of immunofluorescence **(A)** and quantification and statistical analyses. n=50 cells per experiment were counted and the experiment was repeated twice across each genotype (Scale bar, μ100 m). (**B-F**) of mitotic defects observed in wildtype and transgenic MEFs as indicated by the presence of tripolar spindle poles, unaligned metaphase plates, lagging chromosomes, as well as chromatin bridges and micronuclei. Error bars represent the ± SD from two independent experiments. Student’s t-test was performed to determine *P-value* <0.05 (*) and <0.01 (**). **(G)** Representative images of detryosinated (red) and acetylated tubulin of both anaphase and midbody cytokinetic bridges showing stabilized tubulin in *Cep55^Tg/Tg^* MEFs. n=50 cells per experiment were counted and the experiment was repeated twice across each genotype (Scale bar, 100μm). **(H)** Statistical representation showing reduction in mitotic defects (described previously in A-F) upon *KIF2B* overexpression in the MEFs of indicated genotypes. Error bars represent the ± SD from two independent experiments. Student’s t-test was performed to determine *P-value* <0.05 (*) and <0.01 (**).

Increased kinetochore–microtubule (k-MT) stability causes incomplete segregation of DNA, including lagging chromosomes during anaphase^25, 26^. As CEP55 is recruited to centrosomes and spindle microtubules during mitosis^3^ and efficiently bundles microtubules^27^, we asked if *Cep55* stabilizes microtubules, and hence increasing segregation errors during mitosis. To analyse spindle microtubule stability, mitotic cells were stained with antibodies that recognize stable detyrosinated- and acetylated-microtubules. *Cep55^Tg/Tg^* mitotic cells exhibited enhanced detyrosinated- and acetylated-microtubule staining compared to mitotic *Cep55^wt/wt^* cells, indicating these cells have stabilised microtubules in spindle poles and midbodies (Fig 6G). Next, to confirm that the generation of chromosome segregation errors, including lagging chromosomes, in response to *Cep55* overexpression is due to stabilised microtubules, we expressed GFP-tagged *KIF2B*, microtubule depolymerizing kinesin-13 protein, in both *Cep55^Tg/Tg^* and *Cep55^wt/wt^* MEFs. In particular, exogenous expression of *KIF2B* in *Cep55^Tg/Tg^* cells significantly reduced the frequencies of lagging chromosomes, anaphase bridges and micronuclei (Fig 6H), suggesting that overexpression of *Cep55* stabilises microtubules that in part leads to the mitotic defects observed in these MEFs.

### Cep55 overexpression leads to altered Chk1 distribution causing replication stress in an Akt-dependent manner

It has been well established that oncogenes often accelerate DNA replication fork progression and thereby promote GI^28, 29^. We observed that *Cep55*-overexpressing MEFs progressed faster through interphase and entered mitosis more rapidly with enhanced mitotic defects (Fig. 5D and Supp. Fig 7D), including anaphase chromatin-bridge formation that is commonly derived due to replication-associated stress^30^. Since DNA replication is a rate-limiting step during interphase, we therefore investigated the impact of *Cep55* overexpression on replication by examining the replication fork progression rate using DNA fibre assays. We found that the *Cep55-*overexpressing MEFs exhibited a significant increase in replication fork speed (median speed: 1.47kb/min) compared to wildtype cells (median speed: 1.03kb/min) (Fig 7A-B). On the contrary, transient silencing of *Cep55* in these cells significantly reduced replication fork speeds, suggesting that *Cep55* overexpression increases proliferation by allowing cells to replicate faster than the *Cep55^wt/wt^* MEFs (Supp. Fig 9A-B). An increase in fork speed by 40% above the normal fork progression speed can induce DNA damage and genome instability^29^. Next, we investigated the impact of increased replication speed on DNA damage in the *Cep55^Tg/Tg^* MEFs. Interestingly, we initially observed that the *Cep55^Tg/Tg^* MEFs exhibited significantly higher percentage of β-H2AX positive cells (>5 γ-H2AX foci per cell, Supp. Fig 9C) when compared to the *Cep55^wt/wt^* MEFs (Supp. Fig 9D-E). Likewise, we found that *Cep55^Tg/Tg^* MEFs have a higher percentage of EdU positive cells (Fig 7C), compared to *Cep55^wt/wt^* MEFs. Notably, an increase in the percentage of γ-H2AX positive cells was seen in both Edu-positive and Edu-negative population of the Cep55Tg/Tg MEFs, suggesting that DNA damage is persistent. Despite this increase in baseline damage, no significant differences in DNA damage response signalling were apparent between these lines when cells were challenged with 6-Gy γ-irradiation (Fig 7D). However, we noticed a marked reduction in total Chk1 levels in *Cep55^Tg/Tg^* MEFs (Fig 7D). ATR-dependent Chk1 is a well-established effector of DNA damage and replication stress response which is also required for faithful chromosome segregation^31^. Overexpression and/or hyper-activation of AKT has previously been associated with cytoplasmic sequestration of CHK1, hence loss of its checkpoint activity that can ultimately lead to genomic instability^32^. Since *Cep55^Tg/Tg^* MEFs have highly elevated Akt signalling (Fig 3G), we initially investigated the subcellular Chk1 distribution in MEFs of different *Cep55* genotypes. Compared to *Cep55^wt/wt^* MEFs, the *Cep55^Tg/Tg^* MEFs show relatively higher Chk1 levels in cytoplasmic but reduced levels in nuclear fraction (Fig 7E). Meanwhile, treatment of *Cep55^Tg/Tg^* MEFs either with PI3K or AKT inhibitor markedly altered the localisation of Chk1 from cytoplasmic to nuclear fraction, confirming that the activation of Akt signalling in *Cep55*-overexpressing cells sequesters Chk1 in the cytoplasmic fraction. To further confirm the involvement of an Akt-mediated checkpoint defect during replication-mediated stress, we treated *Cep55^Tg/Tg^* MEFs with either BEZ235 or AKTVII inhibitors and performed DNA fibre assay. Our data showed that treatment of *Cep55*-overexpressing cells with Akt inhibitors significantly reduced replication fork speeds compared to DMSO treated cells (Fig 7F and G and Supp. Fig 9F). AKT phosphorylates CHK1 at serine 280 and impairs its nuclear localization and checkpoint activity independent of ATR^32^. To determine the crucial role of Cep55-Akt-dependent checkpoint deficiency, we transiently reconstituted *Cep55^Tg/Tg^* cells with S280A mutant (that cannot be phosphorylated by active-AKT), and *Cep55^wt/wt^* cells with S280E mutant (mimics constitutive AKT-dependent phosphorylation). Our data showed that while S280E mutant significantly increased replication fork speed in *Cep55^wt/wt^* cells, the S280A mutant reconstituted *Cep55^Tg/Tg^* cells on contrary show significantly decreased replication fork speed, suggesting that the checkpoint activity is impaired in Cep55-Akt-dependent manner in these cells (Fig 7H, Supp. Fig). Collectively, our data suggests that overexpression of *Cep55* partially impairs Chk1-mediated checkpoint activation leading to faster replicating cells with persistent DNA damage that undergo an aberrant mitosis, thereby promoting GI in our model.

**Figure 7:**
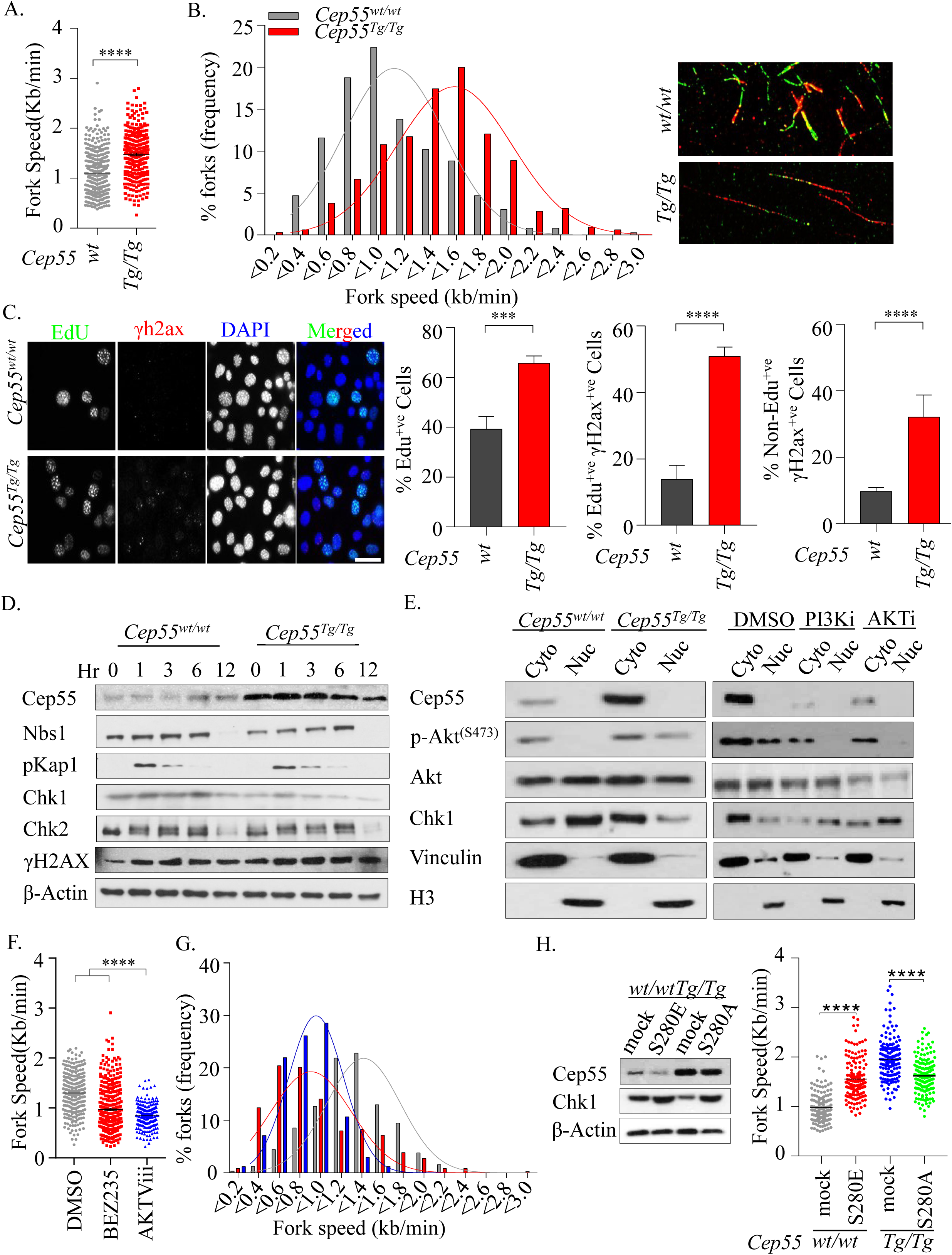
Cep55 overexpression causes replication stress. **(A, B)** Statistical representation of velocity of progressing forks **(A)** and distributions of replication fork speeds **(B)** was determined using DNA fiber analysis. Indicated MEFs were pulsed labeled with IdU (red) and CldU (green) for 25 minutes and the fibers were imaged and quantified. Representative images of respective genotypes are shown on the right hand panel. At least 300 fibers from each cell line were analysed from two independent experiments with error bars representing the standard error of the mean (SEM). Unpaired t test with and without Welch’s correction between two groups was used to determine the statistical *P-value*, <0.0001 (****). **(C)** Representative images of immunofluorescence of EdU (S-phase cells) positivity (green) allowed to label for an hour alongside double stranded marker yH2ax (red) observed in the MEFs of indicated genotypes are shown on the left hand panel. DNA was marked using DAPI (blue). The statistical representation of the percentages of EdU positive cells; γh2ax in EdU positive or negative cells are demonstrated in the right hand side panel. Error bars represent the ± SD from two independent experiments. Student’s t-test was performed to determine *P-value* <0.001 (***) and <0.0001 (****). **(D)** Immunoblot analysis of the whole cell lysate from respective MEFs highlighting the presence of indicated proteins in cells of indicated genotypes after challenged with 6-Gy irradiation. β-Actin was used as loading control. **(E)** Immunoblot analysis of cytoplasmic-nuclear fractionation was performed to determined Chk1 protein distributions with and without indicated inhibitor treatments. Cells were treated for 6 prior to the assay. H3 and Vinculin were used as loading control in each fraction. **(F, G)** Statistical representation of velocity of progressing forks **(F)** and distributions of replication fork speeds **(G)** was determined using DNA fiber analysis as described in A. *Cep55^Tg/Tg^* MEFs were pretreated for 6 hours with indicated inhibitors and forks speeds were determined. Representative images are shown on the right. At least 300 fibers from each cell line were analysed from two independent experiments with error bars representing the standard error of the mean (SEM). One way ANOVA with Brown-Forsythe test was used to determine *P-value* <0.0001 (****). **(H)** Left: Immunoblot analysis showing transiently transfected mutants Chk1 along with Cep55. β-Actin was used as loading control. Right: Statistical representation of velocity of progressing forks as indicted in A. Both cell lines were transiently transfected with 1.5µg indicated mutant constructs respectively (CHK1-S280A and S280E) for 24 h and DNA fiber analysis and immunoblotting were performed. For fiber assays, at least 150 fibers from each cell line were analysed from two independent experiments with error bars representing the standard error of the mean (SEM). One way ANOVA with Brown-Forsythe test was used to determine *P-value* <0.0001 (****).

## Discussion

We previously reported a *Cep55*-overexpression mouse model that exhibits male-specific sterility by suppressing Foxo1 nuclear retention through hyperactivation of Pi3k/Akt signalling^17^. In this study, using the same mouse model, we demonstrate for the first time that *Cep55* overexpression causes spontaneous tumorigenesis. Our data highlights the allele-dependent impact of *Cep55* overexpression on cell proliferation and tumorigenesis *in vivo*. The homozygous *Cep55^Tg/Tg^* mice are prone to develop a wide spectrum of tumours (both solid and haematological origin) with a high incidence rate and high metastasis potential. Interestingly, heterozygous *Cep55^wt/Tg^* mice developed a lower percentage of adenomas (∼20%) and hyperplasia (∼8%), suggesting that single copy overexpression of *Cep55* is sufficient to initiate tumorigenesis, although the latency significantly differs between *Cep55^Tg/Tg^* and *Cep55^wt/Tg^* mice. Notably, the *Cep55^Tg/Tg^* mice demonstrated a higher incidence of lymphomas and sarcomas compared to other type of malignancies, mimicking the phenotype observed in *Trp53^-/-^* mice. As p53 negatively controls CEP55 expression^13^, using a bi-transgenic mouse model, we also demonstrated that single copy overexpression of *Cep55* is sufficient to accelerate heterozygous *Trp53^+/-^* loss*-*induced tumorigenesis. Interestingly, these data also illustrate that either loss or mutation of *Trp53* might be an early event and a critical secondary hit required for tumour initiation observed in the *Cep55^Tg/Tg^* mice. Consistent with this, we observed high p53 protein levels, which are most likely an indication of mutated *Trp53*, in representative *Cep55^Tg/Tg^* tumour tissues than normal adjacent tissues. Notably, partial depletion of *Cep55* (50%) in TCLs significantly delayed tumour initiation and progression, while near-complete depletion (90%) totally impaired tumour initiation in a xenograft model.

As *Cep55* has been linked with GI and its overexpression causes a wide range of tumours *in vivo*, we further characterised GI in *Cep55*-overexpressing cells. *Cep55^Tg/Tg^* MEFs exhibited a high level of cytokinesis failure accompanied by genome doubling. Importantly the *Cep55^Tg/Tg^* MEFs showed high level of numerical and structural CIN compared to MEFs of other genotypes. Importantly, in this study, we showed for the first time that *Cep55* overexpression causes mitotic defects including a high frequency of chromatin bridge and micronuclei formation during anaphase. As CEP55 is a microtubule-bundling protein^27^, missegregation of chromosomes upon *Cep55* overexpression might be indicative of kinetochore-microtubule (k-MT) hyperstability. Consistent with this notion, we found that overexpression of *Cep55* stabilised microtubules and predisposed cells to CIN. Notably, reducing microtubule stability by forced expression of KIF2b in *Cep55^Tg/Tg^* MEFs significantly reduced lagging chromosomes. The influence of *Cep55* overexpression on sister chromatid segregation errors accompanied by cytokinesis failure explains the delayed mitotic exit observed in the *Cep55*-overexpressing cells. Taken together, our data suggests that hyperstabilised microtubules and defective cytokinesis in Cep55-overexpressing cells might be major source of chromosome segregation errors and tetraplodization that can predispose these cells to genomic instability which over time might facilitate tumour development.

Consistent with previous reports (reviewed by Jeffery *et al*. ^4^), *Cep55* overexpression led to rapid proliferation. We observed that the *Cep55^Tg/Tg^* MEFs displayed hyper-phosphorylated Akt and deregulated downstream PI3K/ Akt signalling such as Gsk-3β, Myc and β-Catenin which might be a further source of genomic instability in these cells. Akt hyperactivation is known to result in cytoplasmic sequestration of Chk1, this might result in a compromised S-phase checkpoint that increases replication fork progression in *Cep55^Tg/Tg^* MEFs to allow uncontrolled cell cycle progression and consequently promote genomic instability. Consistent with this, overexpression of CHK1 mutant (S280A), that cannot be phosphorylated by overactive AKT, in *Cep55^Tg/Tg^* MEFs or their treatment with PI3K/AKT pathway inhibitors resulted in reduced fork-progression. Furthermore, loss of Chk1 function has also been shown to induce chromosomal segregation errors and chromatin-bridges during anaphase resulting in CIN^31, 33^, resembling the phenotype we observe.

Deregulation of mitotic proteins have long been known to contribute to early cellular transformation and tumorigenesis^34^ though they are rarely mutated in cancer ^35, 36^, but rather prone to amplification. Abnormal expression (loss or gain) of critical mitotic proteins, especially those included in the CIN70 gene signature, such as *MAD2*^37^, *BUB1*^38^, *AURKA*^39^, *EMI1*^40^, PLK1^41, 42^, *TTK1*^43^ and many more, at the genetic level have been shown to induce spontaneous tumorigenesis. The major phenotype observed in these mouse models was defective chromosomal segregation during anaphase which led to CIN and genomically unstable malignancies, similar to the phenotype observed in our model. Thus, the interplay of these mitotic genes with *Cep55* overexpression needs further evaluation. Importantly, in our previous study in breast cancer, we have shown that CEP55 overexpression protects aneuploid cells during perturbed mitosis^17^. We have demonstrated that high level of CEP55 significantly induced mitotic slippage in TNBCs as loss of CEP55 enables mitotic cell death by enabling premature mitotic entry upon being challenged with anti-mitotic drugs. Consistently, herein we have demonstrated that Cep55 is a protector of aneuploidy during aberrant mitosis as the aneuploid *Cep55^Tg/Tg^* MEFs underwent mitotic slippage in response to anti-mitotic drugs and survived mitotic cell death. It also explains the ability of the highly polyploid *Cep55^Tg/Tg^* MEFs to re-enter mitosis and continue proliferation as Cep55 overexpression allows high tolerance and better survival of these cell populations.

A recent report has suggested that cells procure specific genomic alterations, mainly impacting the regular function of mitotic genes prior to malignant transformation^44^. CEP55 overexpression has been linked with tumorigenesis for a wide-variety of cancers. However, this is the first report to our knowledge demonstrating that overexpression of Cep55 has a causative role in tumorigenesis. Our data clearly demonstrates that Cep55 overexpression beyond a critical level is self-sufficient to induce a wide spectrum of spontaneous tumours. Importantly, we have shown that *Cep55* overexpression leads to induction of pleotropic events such as PI3K/Akt pathway activation, Chk1 sequestration compromising the replication checkpoint, and stabilized microtubules along with chromosomal segregation anomalies which all together causes CIN. Accumulation of these anomalies over time might induce tumourigenesis. In summary, our mouse model could be a valuable tool in studying the mechanism of CIN-associated tumorigenesis and development of CIN-targeting therapies.

## Materials and Methods

### Reagents

Nocodozole, BEZ235, BKM120, AZD6244 and AKTVIII were purchased from Selleck Chemicals LCC. Small interfering RNAs (siRNAs) were from Shanghai Gene Pharma. Dulbecco’s Modified Eagle’s Media (DMEM), Click-iT Alexa Fluor 488 EdU (5-ethynyl-2’-deoxyuridine) imaging kit and Lipofectamine RNAiMAX was purchased from Life Technologies. Foetal Bovine Serum (FBS) was purchased from SAFC Biosciences™, Lenexa, USA. CellTiter 96^®^ AQueous One Solution Cell Proliferation Assay and Dual-Glo® Luciferase Assay were purchased from Promega Corporation.

### Animal husbandry and ethics statement

All animal work was approved by the QIMR Berghofer Medical Research Institute, Animal Ethics Committee (number A0707-606M) and was performed in strict accordance with the Australian code for the care and use of animals for scientific purposes. The animals were maintained as per the guidelines reported previously^17^.

### Histopathological analysis and immunohistochemistry

For histologic examination, tissues were collected and fixed in 4% formaldehyde in PBS as per the standard protocol described previously^17^.

### Cell Culture and synchronization

To generate the MEFs, mice pregnancy was accessed on the basis of a copulation plug on the following morning post mating date, designated as embryonic day. Such assessment was done for isolating MEFs E13.5 and single cell isolation was performed using the standard protocol described previously^45^. To generate the primary tumor lines (TCLs), tumor was surgically removed followed by mechanical disaggregation using a sterile scalpel blade and then incubation in 0.1% collagenase (Sigma Aldrich) in 10 mL of DMEM containing 20% FBS and 1% penicillin-streptomycin (100 U/mL), 1% L-glutamine and cultured in a 25 cm^2^ tissue flasks. After 24 hrs, the cells were trypsinized and cultured in a new 25 cm^2^ tissue flask with media supplemented with 100 µL (100 µg/mL) of EGF, 500 µL (10mg/mL) of insulin and 1% Sodium pyruvate (Life Technology^TM^. The culture of the murine cell lines was maintained by incubating at 37 °C with 20% oxygen levels and 5% CO_2_. Cells were synchronized at G1/S by double thymidine block as described previously^46^.

### Genotype analysis and Quantitative real-time PCR

Genotyping, RNA extraction and quantitative real-time PCR was performed using the primer sets used in these assays were used as per the protocol described previously^17^.

### Immunoblot analysis

The protein extraction from cell lysate or tissue lysate were prepared in urea lysis buffer (8M urea, 1% SDS, 100mM NaCl, 10mM Tris (pH 7.5) and incubated for 30 minutes on ice after which the samples were sonicated for 10 seconds. Western blotting was performed as per the standard protocol and some of the antibodies used for immunoblotting has been described previously^17^. The following are additional antibodies used in this study: γH2AX S139 (05-636); Cell Signaling antibodies: PARP (#9542), pAKT^S473^ (#4060), AKT (#9272), pPdk1^S241^ (#3061), Pdk1(#3062) Chk1 (2G1D5) (#2360), p-GSK-3β^(Ser9)^ (#9336), GSK-3β (#9315), p-Histone H3 (#9706); Millipore antibody: Chk2 (Clone 7) (05-649); BD Pharmingen antibody: β-actin (612656); Bethyl antibody:pKap1^(S824)^ (A300-767A).

### Cell proliferation assay

The cell proliferation assay using The IncuCyte® S3 Live-Cell Analysis system (Essen BioSciences Inc, USA), as described previously^18^. Doubling time was analyzed at every 12 hour interval by counting the overall cell population compared to originally seeded population using the Countess® automated cell counter (Life Technologies^TM^). The NIH-3T3 proliferation assay was per performed by using the standard protocol as described previously^45^.

### Colony formation assays

Five hundred to one thousand cells were seeded on 12 well plates and incubated for additional 14 days to determine colony viability. The colonies were fixed with 0.05% crystal violet for 30 minutes as described previously^18^.

### Flow cytometry and cell cycle analysis

Cell cycle perturbations and the subG1 apoptotic fractions were determined using flow cytometry analysis of cells stained with propidium iodide and analyzed using ModFit LT 4.0 software as described previously^18^.

### Immunofluorescence

Cells were seeded and incubated overnight on coverslips and were fixed for 15 minutes in 4% paraformaldehyde in PBS, permeabilized in 0.5% Triton X-100-PBS for 15 minutes and blocked in 3% filtered bovine serum albumin (BSA) in PBS. Coverslips with primary antibodies were diluted in blocking solution and incubated overnight at 4°C. Alexafluor conjugated secondary antibodies were diluted 1/300 and DAPI (diluted 1/500 in blocking buffer, stock 1mg/ml), in blocking solution and stained for 45 minutes at 37°C in humidifier chamber. Slides were washed thrice with 0.05% Tween 20 in PBS and mounted in Prolong Gold. Slides were imaged using GE DeltaVision Deconvolution microscope and analyzed using Image J as described previously^18^. Antibodies used for immunofluorescence were: γH2AX S139 (05-636; Millipore), p-Histone H3 (#9706; CST), α-Tubulin (T9026) and γ-Tubulin (T5192).

### DNA Combing Assay

The DNA fiber protocol was followed as described previously us and others^47, 48^. Cells were labelled with CldU and IdU for 20 minutes each. Progressive replication fork speed was calculated based on the length of the CldU tracks measured using ImageJ software. At least 300 replication tracks were analyzed for each sample in two independent experiments. The fork speed was calculated based on conversion factor 1 µm=2.59kb^49^.

### Gene silencing

Transient gene silencing was performed by reverse transfection using 10 nM of respective small interfering RNAs (siRNAs). The sequences involved *Cep55*_Scr (5’CAAUGUUGAUUUGGUGUCUGCA3’); *Cep55*_SEQ1 (5’ CCAUCACAGAGCAGCCAUUCCCACT 3’) and *Cep55*_SEQ2 (5’ AGCUACUGAGCAGUAAGCAAACAUU). The siRNAs were manufactured by Shanghai Gene Pharma. The transfection was performed using Lipofectamine RNAiMAX (Life Technologies^TM^). Mouse small hairpin RNAs (shRNAs) for Cep55 (pLKO plasmids, (Sigma Aldrich®, St Louis, USA)) clones were established using lentiviral packaging using PEI (Poly -ethyleneimine) solution (Sigma Aldrich®, St Louis, USA). *Cep55*_Scr (5’CCGGCGCTGTTCTAATGACTAGCATCTCGAGATGCTAGTCATTAGAACAGCGT TTTTT3’); *Cep55*_sh#1 (5’CCGGCCGTGACTCAGTTGCGTTTAGCTCGAGCTAAACGCAACTGAGTCACGGT TTTTG); *Cep55*_sh#2 (5’CCGGCAGCGAGAGGCCTACGTTAAACTCGAGTTTAACGTAGGCCTCTCGCTGT TTTTG3’); *Cep55*_sh#3 (5’CCGGCGTTTAGAACTCGATGAATTTCTCGAGAAATTCATCGAGTTCTAAACGT TTTTT3’); *Cep55*_sh#4 (5’CCGGGAAGATTGAATCAGAAGGTTACTCGAGTAACCTTCTGATTCAATCTTCT TTTTT3’).

### Live cell imaging

Live cell imaging for double thymidine releases was performed on an Olympus IX81 microscope using excellence rt v2.0 software. Images were analyzed using analySIS LS Research, version 2.2 (Applied Precision) as described previously^50^. Live cell imaging for tracking mitotic defects was performed in H2B Cherry transfected MEFs of each genotype using 20X Andor Revolution WD - Spinning Disk microscope.

### In vivo xenografts

All mice were housed in standard condition with a 12h light/dark cycle and free access to food and water. 2.5 x 10^6^ TLC were prepared in 50% matrigel (BD, Biosciences, Bedford, USA)/PBS and injected subcutaneously injected into the right flank of 6 week old NOD/SCID mice as described previously^18^.

### Bioinformatics analysis

Whole-chromosome (WC) and chromosome arm-level (CAL) somatic copy number aberrations (SCNAs) were inferred from TCGA processed (Level 3) Affymetrix Genome Wide SNP6.0 Array data for the indicated cancer types, as previously described ^51^. Using the same datasets, ASCAT2.4 ^52^ was used to compute the ploidy level for each sample. Samples with ploidy between 1.9 and 2.1 were considered diploid, samples with ploidy lower than 1.9 or between 2.1 and 2.5 were called near-diploid aneuploid and samples with ploidy>2.5 were considered aneuploid and having undergone at least one whole-genome doubling (WGD).

### Statistical analysis

Student’s t-test; one-way or two-way ANOVA; RPKM and RSEM with Bonferoni *post hoc* or Mann-Whitney *U* test testing (specified in figure legend) and Fisher exact test was performed using GRAPHPAD PRISM v6.0 (GraphPAd Software, LaJolla, CA, USA) and the p-values were calculated as indicated in figure legends. Asterisks indicate significant difference (*p<0.05, **p<0.01, ***p<0.001 and ****p<0.0001), ns= not significant.

## Acknowledgments

We thank the members of the Khanna laboratory for technical assistance, Stephen Miles for maintaining cell lines, QIMR Berghofer Flow Cytometry and Animal facility staffs, Nigel Waterhouse and Tam Hong Nguyen from ACRF Imaging Centre for microscopic assistance, and Paul Collins for STR profiling and Mycoplasma testing. We thank Professor Rajiv Khanna for DS salary supports and Dr. Hiroyuki Niida from Hamamatsu University School of Medicine for providing mutant Chk1 constructs.

## Author Contributions

Conceptualization, DS, MK and KKK; Investigation and data analysis, DS, PN. DN, AB, PR, VAJS, ALB, GS, MW, JWF, and MK; Bioinformatics, PHGD; Writing–Original Draft, DS, MK, and KKK; Writing–Review & Editing, all authors. All authors read and approved the final manuscript.

## Supplementary figure legends

***Supp. Fig1: CEP55 is overexpressed in a broad range of cancers independent of a proliferation-associated effect.***

**(A)** CEP55 expression in multiple data sets of various cancer types, analyzed using the Oncomine database^53^. Overall, 7403 tumor samples to 1467 normal control samples of matched tissue type were compared and we observed that the expression of *CEP55* to be significantly higher than in matched normal tissue. A total of 212 data sets were identified showing statistically significant deregulated expression of CEP55 in tumours compared to matched normal control tissues. Each box represents a dataset. Studies showing significant CEP55 overexpression are shown in red, those showing significant underexpression in blue. Fold over- or under-expression is shown as indicated.

**(B)** Pie charts derived from the dataset described in (A) implying significant CEP55 expression differences compared to normal tissue (n=212) wherein 193 case studies (91%) illustrated significant overexpression, while 19 case studies (9%) showed significant underexpression. Proportion of studies showing significant overexpression (red) and underexpression (blue) is shown. Fisher’s exact tests was used to determine the respective *P*-values.

**(C)** RSEM-normalised CEP55 expression levels, performed using the TCGA^54^ dataset, in lung adenocarcinoma (LUAD), liver hepatocellular carcinoma (LIHC) and or colorectal adenocarcinoma (COADREAD) samples compared to matched normal control samples. Collectively, the datasets illustrate that *CEP55* is significantly upregulated in respective tumors in comparison to normal control tissue. Student t’ tests was used to determine *P-value* <0.0001 (****).

**(D)** Ratios of CEP55/MKI67 (upper panel) and CEP55/PCNA (lower panel) RSEM-normalised expression levels in LUAD, LIHC and COADREAD of datasets as in (C).

**(E)** Representation of the tumors that express MKI67 at levels in the same range as normal samples show significantly elevated CEP55 expression levels than the normal samples. These data illustrate that expression of CEP55 is significantly higher in tumors than in normal tissue, even after compensation for the expression of the cell proliferation markers Ki67 (upper panel) or PCNA (lower panel), indicating that CEP55 expression in tumors is cell cycle-independent. Mann-Whitney *t-* test was used to determine *P-value* <0.0001(****).

***Supp. Fig2:* Spontaneous tumourigenesis induced by *Cep55* overexpression *in vivo*.**

**(A)** Statistical representation of the body weight observed at the end of tumor survival of each genotype (n>40 per group). Error bars represent the ± SEM from the entire experimental cohort. One-way ANOVA test was performed to determine *P-value* <0.0001 (****).

**(B)** Representation of H&E-stained microscopic images indicated tumor lesions from different organs of tumors-bearing *Cep55^Tg/Tg^* mice; (scale bars, 200µm).

**(C-F)** Statistical representation of the overall distribution of indicated tumour lesions among respective major organs of tumour-bearing *Cep55^Tg/Tg^* mice (HCC=hepatocellular carcinoma). Percentage of tumor burden **(C)**, types of tumors that they originated from **(D)**, Sarcoma **(E)** and hyperplasia **(F)** observed in tumour-bearing *Cep55^Tg/Tg^* mice.

**(G)** Statistical representation of the respective weights of spleen, liver and lung observed in the indicated genotypes at the end of their respective survival. Error bars represent the ± SEM from the entire experimental cohort. One-way ANOVA test was performed to determine *P-value* <0.0001 (****).

**(H)** Percentage of metastasis incidence observed in the tumor bearing *Cep55^Tg/Tg^* mice.

***Supp. Fig3:* Loss of Trp53 leads to early tumor latency in *Cep55* overexpressing mice.**

**(A)** Boxplots showing CEP55 expression in indicated tumors or matched normal samples with *TP53* copy number and/or mutation status as indicated. Numbers of samples for each column are shown above the x-axis. *P*-values were determined using Mann-Whitney *t-* test. ****p<0.0001.

**(B)** Representation of H&E stained microscopic images of selected sections of indicated tumors lesions from different organs of tumor-bearing mice of respective genotypes; (scale bars, 200µm).

**(C)** Percentage of the overall distribution of indicated tumor lesions among respective major organs of tumor-bearing mice of indicated genotypes.

**(D)** Percentage of number of tumors (tumour burden) observed in each mice of respective genotypes.

**(E)** Statistical representation of the body weight observed at the end of tumor survival of each genotype (n 10 per group). Error bars represent the ± SEM from the entire experimental ≥ cohort. One-way ANOVA test was performed to determine *P-value* <0.0001 (****).

**(F)** Statistical representation of the respective weights of spleen, liver and lung observed in the indicated genotypes at the end of their respective survival. Error bars represent the ± SEM from the entire experimental cohort. One-way ANOVA test was performed to determine *P-value* <0.0001 (****).

***Supp. Fig4:* Cep55 overexpression promotes cell proliferation advantage *in vivo*.**

**(A)** Representation of genotyping of DNA isolated from the primary MEFs of each indicated genotype using PCR, showing the presence of amplicons of the expected size for each genotype. #1 and #2 denotes biologically independent DNA samples of each genotype.

**(B)** Statistical representation of relative fold change in the doubling time of immortalized MEFs from each genotype as indicated in (C) (n=3 per group). Error bars represent the ± SEM from the entire experimental cohort. One-way ANOVA test was performed to determine *P-value* <0.001.

**(C)** Statistical representation of cell proliferation observed in the immortalized MEFs of indicated genotype at different time points during serum starved conditions (n=3 per experiment). One-way ANOVA test was performed to determine *P-value* <0.0001.

**(D)** Immunoblot analysis of indicated whole tissue lysates of respective genotypes of six-month old littermates. β-Actin was used as loading control.

**(E)** Statistical representation of the cell viability of the immortalized MEFs of each genotype after 48hrs of treatment with indicated small molecule inhibitors treated as per the designated concentration. (n=2 per group). Error bars represent the ± SEM from the entire experimental cohort. One-way ANOVA test was performed to determine *P-value* not significant (ns), <0.05 (*) and <0.01 (**).

**(F)** Representative images of cell morphology (bright field image; fluorescence image wherein α-tubulin (green) marks the cytoplasm while DAPI (blue) marks the nucleus) of the TCLs isolated from the haemangiosarcoma found in (Fig1 Ci). The red arrow indicated presence of multinucleated cells while the white arrow represents presence binucleated cells.

**(G)** Immunoblot analysis of the whole cell lysate of TCLs to assess the levels of indicated signaling molecules following transient depletion of *Cep55* using *siCep55* (10 nM, S1). β-Actin was used as loading control (left panel). Effect of Cep55 depletion using siCep55 (10 nM) on cell proliferation, assessed using the IncuCyte ZOOM^®^ live-cell imager. The percentage of cell confluence was determined using an IncuCyte mask analyser (right panel). Error bars represent the ± SD from two independent experiments. Student t’ test was performed to determine *P-value* <0.01 (**).

**(H)** Immunoblot analysis of whole-cell lysates of TCLs to validate extent of *Cep55* depletion with indicated sh*Cep55* sequences (indicated as #S1-#S4 with sh_Scr as control) in the TCLs. β-Actin was used as loading control.

**(I)** Effect of *Cep55* depletion on cell proliferation in TCLs assessed as described in (G). Error bars represent the ± SD from two independent experiments. Student t’ test was performed to determine *P-value* <0.01 (**).

**(J)** Statistical representation of the cell cycle profile of respective TCL clones (n=2 per group).

**(K)** Statistical representation of the cell viability of the indicated TCL clones after 48hrs of treatment with indicated small molecule inhibitors treated as per the designated concentration. (n=2 per group). Error bars represent the ± SEM from the entire experimental cohort. One-way ANOVA test was performed to determine *P-value* not significant (ns), <0.05 (*) and <0.01 (**).

***Supp. Fig5: Cep55 overexpression causes genomic instability.***

**(A)** Boxplots showing CEP55 expression in tumors whose genomes are diploid, near-diploid aneuploid or aneuploid after whole-genome doubling (WGD). Data are from the TCGA liver hepatocellular carcinoma (LIHC), lung squamous cell carcinoma (LUSC), lung adenocarcinoma (LUAD) and colorectal adenocarcinoma (COADREAD) datasets. Mann-Whitney *U* tests was used to determine *P-value* <0.05 (*), <0.01 (**), <0.001 (***), <0.0001 (****).

**(B)** Statistical representation of polyploidy analysis (>4N DNA content) determined using FACS in the respective primary MEFs of each genotype. Error bars represent the ± SD from the entire experimental cohort. One-way ANOVA test was performed to determine *P-value* <0.001 (***).

**(C)** Statistical representation showing comparison of overall polyploidy analysis (>4N DNA content) between primary and immortalized *Cep55^Tg/Tg^* MEFs. Error bars represent the ± SD from the entire experimental cohort. One-way ANOVA test was performed to determine *P-value* <0.001 (***).

**(D)** Statistical representation of overall polyploidy analysis observed in the indicated tissues of respective age-matched mice of each genotype. Error bars represent the ± SD from the entire experimental cohort. One-way ANOVA test was performed to determine *P-value* <0.0001 (****).

**(E)** Immunofluorescence showing genomic instability observed among the respective sh*Cep55*-depleted isogenic clones as indicated by the presence of multiple nuclei (marked by DAPI staining) compared to counterparts. The entire cell (cytoplasm) was marked by α-tubulin (green), while the centrosomes were marked by γ-tubulin (red) (Scale bar, 100μm).

***Supp. Fig6: Association of CEP55 overexpression with aneuploidy.***

**(A)** Boxplots representation showing CEP55 expression in indicated tumors with whole-chromosome (WC)-euploid and WC-aneuploid genomes. The data was defined using the TCGA LIHC, LUSC, and LUAD datasets (described in Supp. Fig5A)^51^.

**(B)** Boxplots as in (A) but at the chromosome arm level (CAL).

**(C)** Boxplots representation demonstrating the chromosome arm-level (CAL) aneuploidy, i.e., total number of chromosome arms gained or lost per sample, with respect to the highest (hi) and lowest (lo) CEP55 mRNA expression quartiles from TCGA RNAseq data.

**(D)** Boxplots representation demonstrating the whole-chromosome (WC) aneuploidy, i.e., the total number of whole chromosomes gained or lost per sample, with respect to the highest (hi) and lowest (lo) CEP55 mRNA expression quartiles from TCGA RNAseq data. For all of the above, Mann-Whitney U tests was used to determine *P-value*.

***Supp. Fig7:* Mitotic cell fate in Cep55 overexpressing MEFs.**

**(A)** Representative images of immunofluorescence demonstrating mitotically active cells observed in *Cep55^Tg/Tg^* MEFs as compared to other counterparts. Mitotic cells are marked by phospho-histone H3 (green) and the nucleus is marked by DAPI (blue). Scale bar, 100 μm (left panel). Statistical representation of phospho-histone H3^+ve^ cells in the MEFs of indicated genotypes (right panel). Error bars represent the ± SD from the entire experimental cohort. One-way ANOVA test was performed to determine *P-value* not significant (ns), <0.001 (***).

**(B)** Statistical representation of mitotic index (number of rounded cells/overall cells in an area) observed in the MEFs of indicated genotypes using bright field Olympus Xcellence IX81 time-lapse microscopy per-field. Overall, 300 cells were counted (∼40 cells per field) of each genotype. Error bars represent the ± SD from the entire experimental cohort. One-way ANOVA test was performed to determine *P-value* not significant (ns), <0.01 (**).

**(C)** Statistical representation of phospho-histone H3^+ve^ cells observed in the respective sh*Cep55* depleted isogenic clones. Error bars represent the ± SD from the entire experimental cohort. One-way ANOVA test was performed to determine *P-value* not significant (ns), <0.001 (***).

**(D)** Data showing the cell cycles profiles of MEFs of indicated genotype. The cells were first synchronized by double-thymidine block and released in regular culture media following which, they were collected after 2-hour intervals. Error bars represent the ± SD from the entire experimental cohort.

**(E)** Statistical representation showing the percentage of binucleated (left panel) and multinucleated cells (right panel) observed in the respective MEFs of indicated genotypes calculated using time-lapse microscopy (n=100 cells of each genotype). Error bars represent the ± SD from the entire experimental cohort. One-way ANOVA test was performed to determine *P-value* <0.05 (*), <0.01 (**), <0.001 (***).

**(F)** Statistical representation of the cell cycle profile of the respective sh*Cep55* depleted isogenic clones in the presence or absence of *nocodazole (0.5* μ*M)* (n=2 per group).

**(G)** Statistical representation of polyploidy analysis (>4N DNA contents) determined using FACS in the respective sh*Cep55* depleted isogenic clones in presence or absence of *nocodazole (0.5* μ*M).* Error bars represent the ± SD from the entire experimental cohort. One-way ANOVA test was performed to determine *P-value* <0.0001 (****).

**(H)** Statistical representation of percentage SubG1 population was determined using FACS in the respective sh*Cep55* depleted isogenic clones in presence or absence of *nocodazole (0.5* μ*M)* Error bars represent the ± SD from the entire experimental cohort. One-way ANOVA test was performed to determine *P-value* <0.0001 (****).

***Supp. Fig8: Cep55 overexpression causes mitotic defects.***

**(A, B)** Representative images showing normal **(A)** and perturbed mitoses **(B)**. Individual cells were tracked using bright-field Olympus Xcellence IX81 time-lapse microscopy and mitotic anomalies were determined (Scale bar, 100μm).

***Supp. Fig9: Cep55 overexpression causes replication stress.***

Statistical representation of velocity of progressing forks **(A)** and distributions of replication fork speeds **(B)** was determined using DNA fiber analysis upon *Cep55* knockdown in *Cep55^Tg/Tg^* MEFs. At least 300 fibers from each cell line were analysed from two independent experiments with error bars representing the standard error of the mean (SEM). Unpaired t test with and without Welch’s correction between two groups was used to determine the statistical *P-value*, <0.0001 (****).

**(C)** Representative images of immunofluorescence (left panel) demonstrating presence of DNA damage marked by γH2ax (green) observed in indicated genotypes (Scale bar, 100μ Statistical representation showing percentage of γH2ax positive cells (>5 foci of yH2ax/cell) in the MEFs of indicated genotypes MEFs (right panel). Error bars represent the ± SD from the entire experimental cohort. One-way ANOVA test was performed to determine *P-value* <0.01 (**).

**(E)** Immunoblot analysis of indicated proteins in cells challenged with 6-Gy irradiation. β-actin was used as loading control.

**Supp. Table 1:**
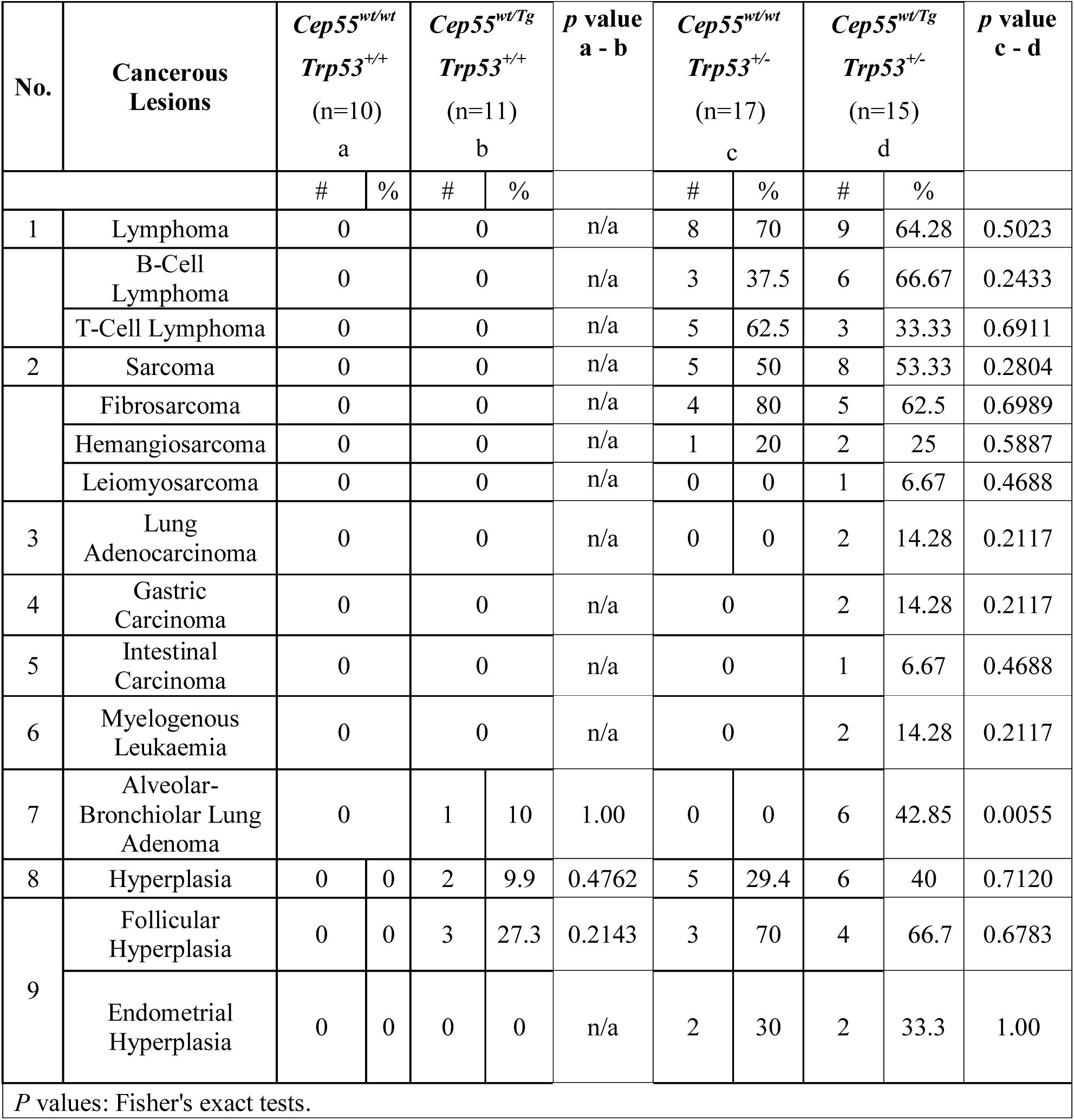
Distribution of cancer spectrum in bi-transgenic mice.

