## Supplementary figures and images for "Cep55 overexpression promotes genomic instability and tumorigenesis in mice"

### Supp. Figures

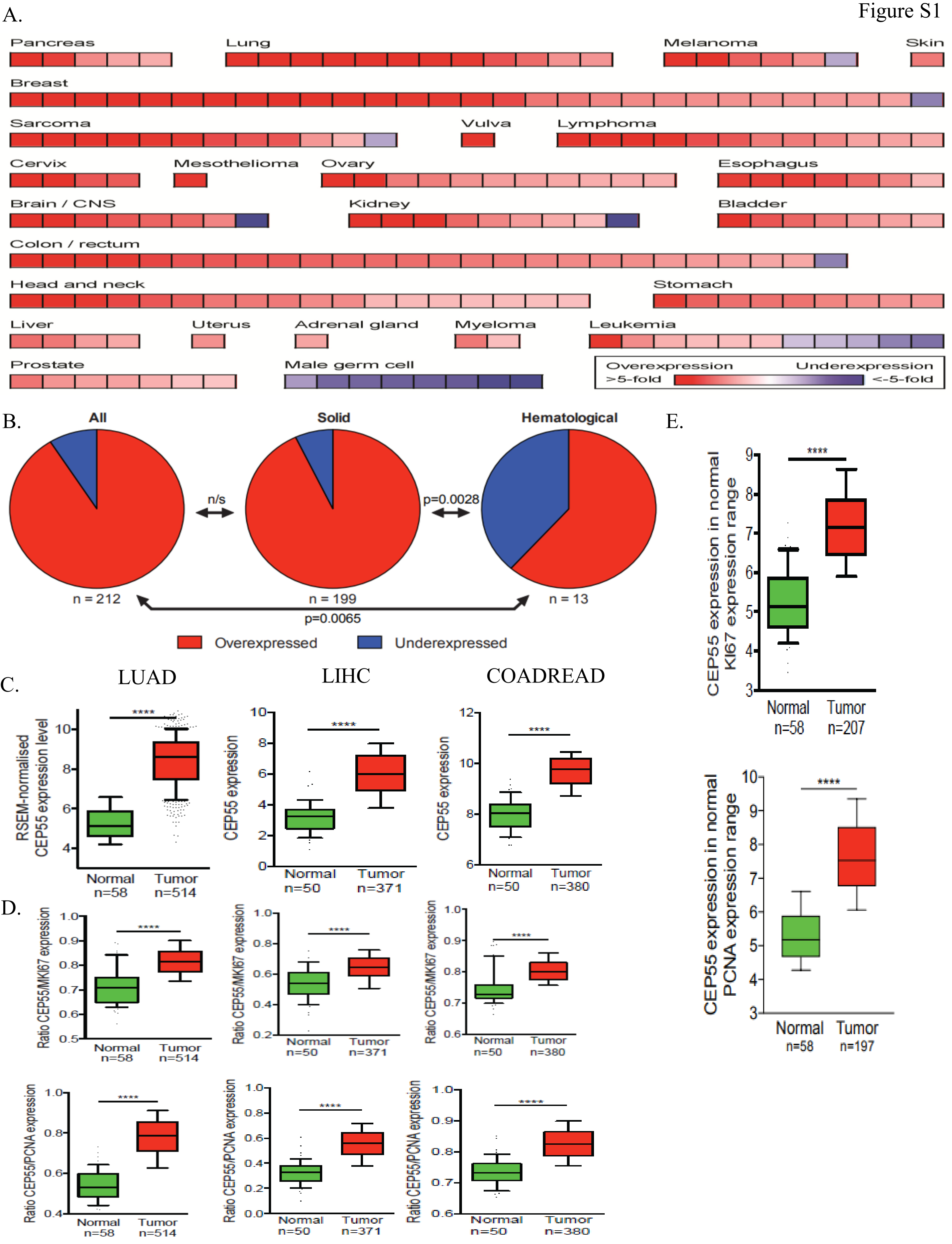

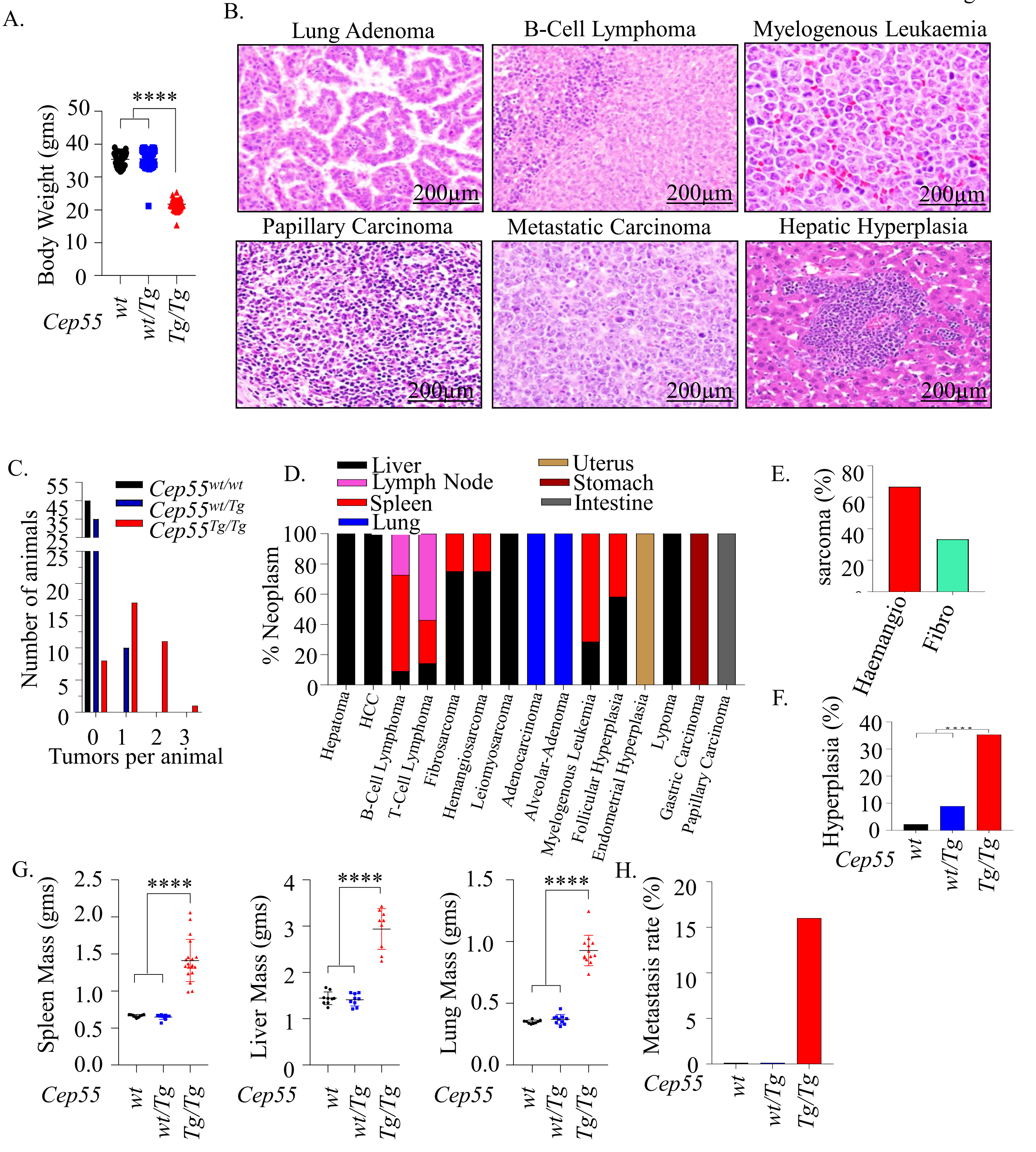

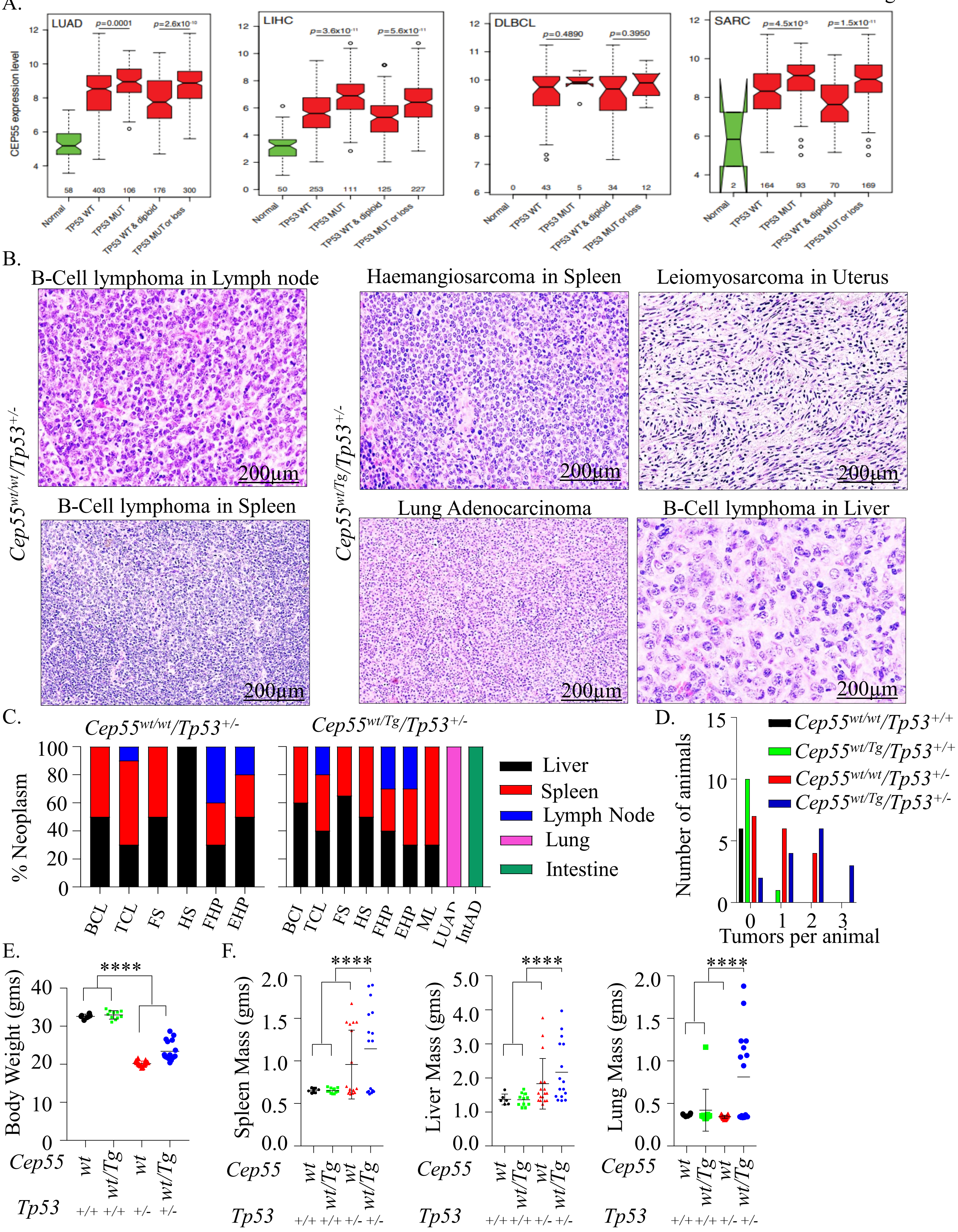

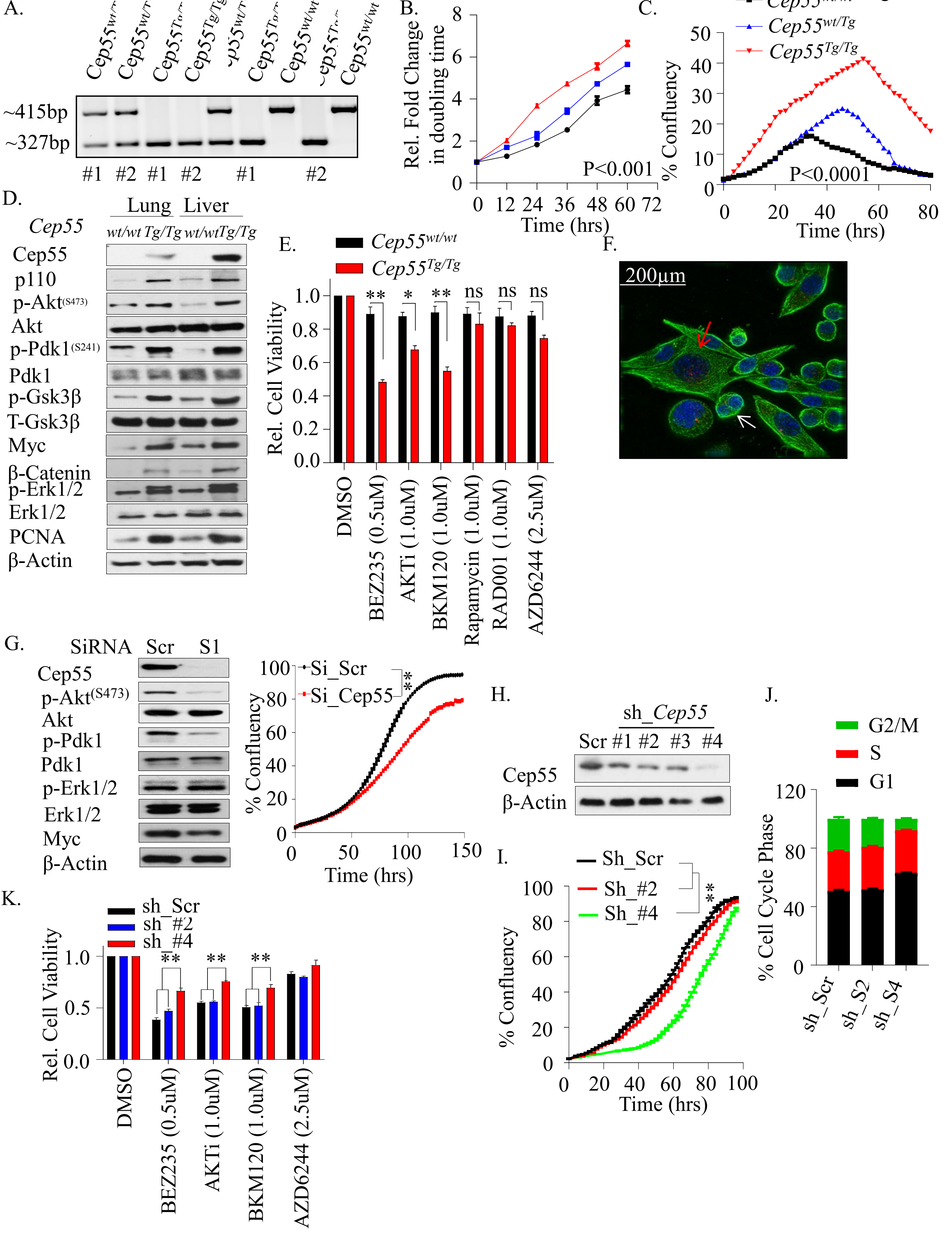

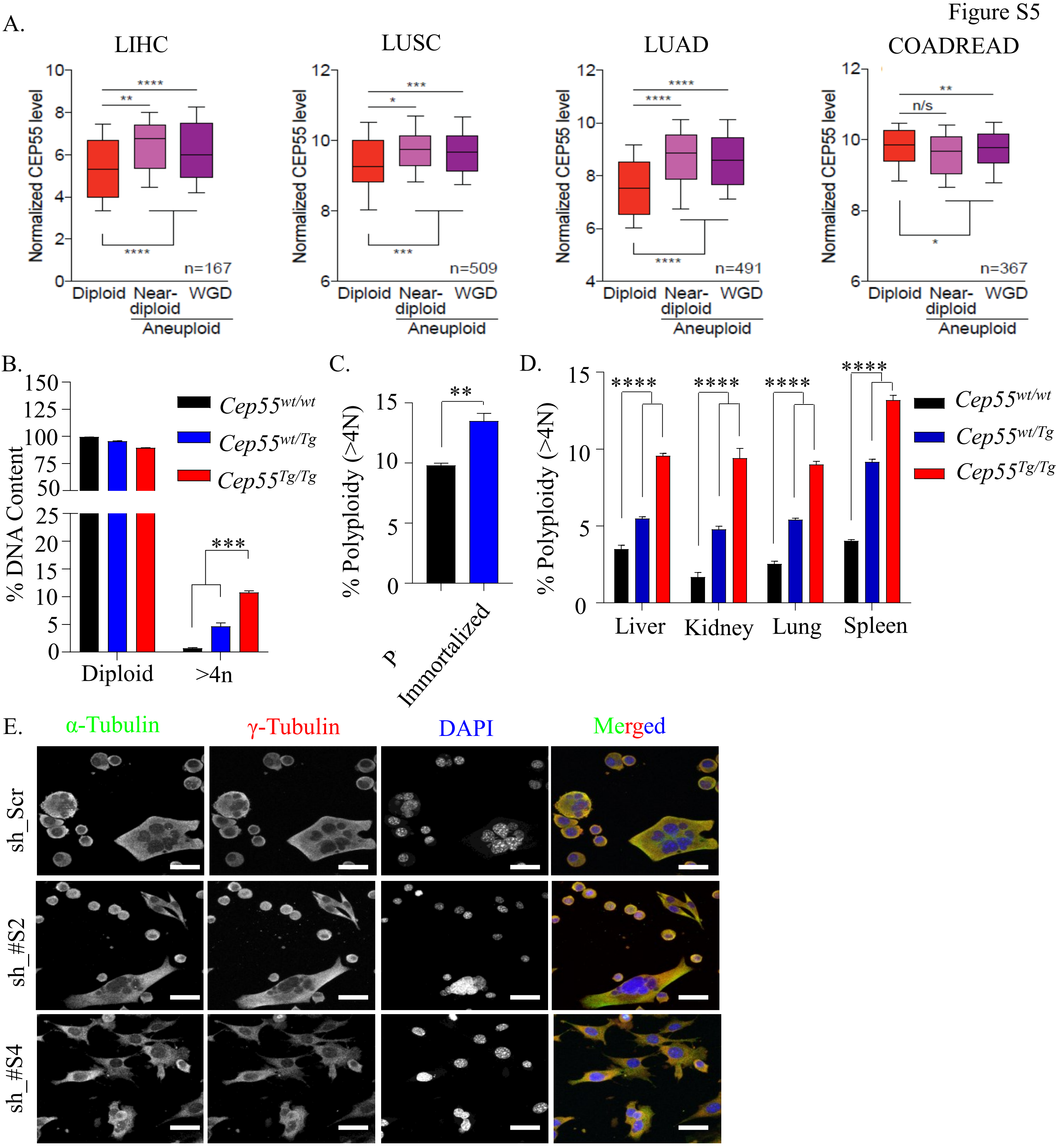

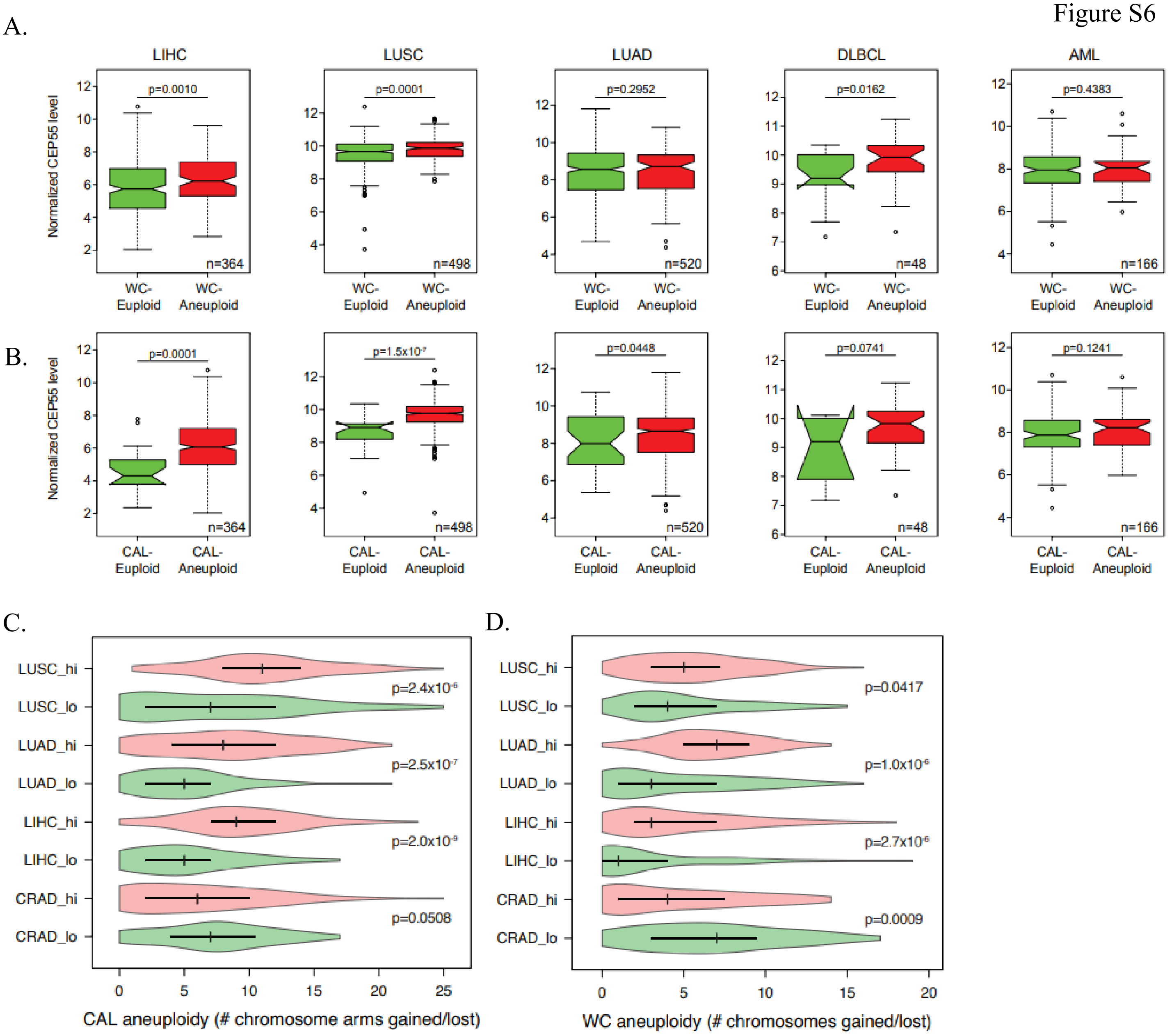

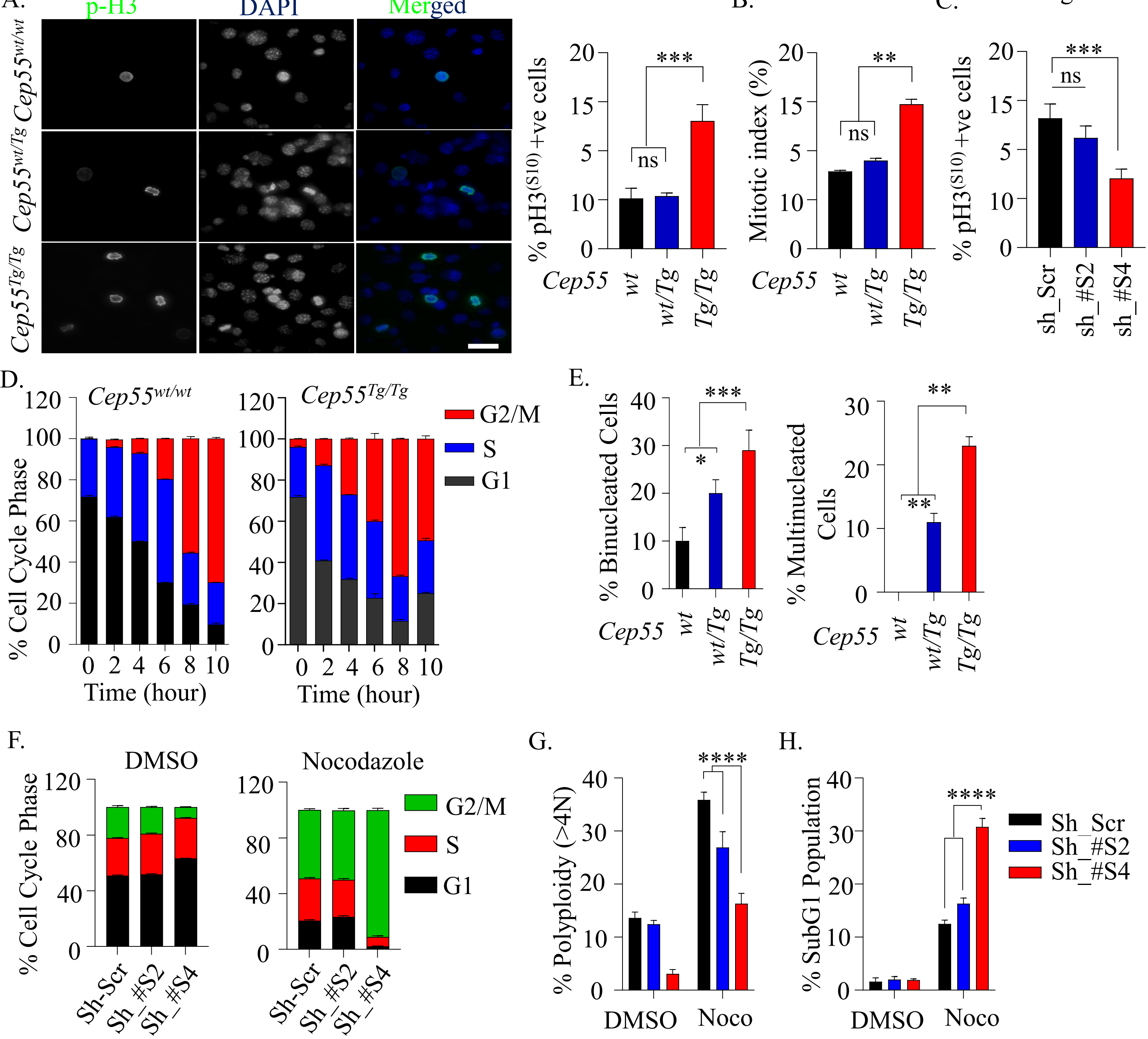

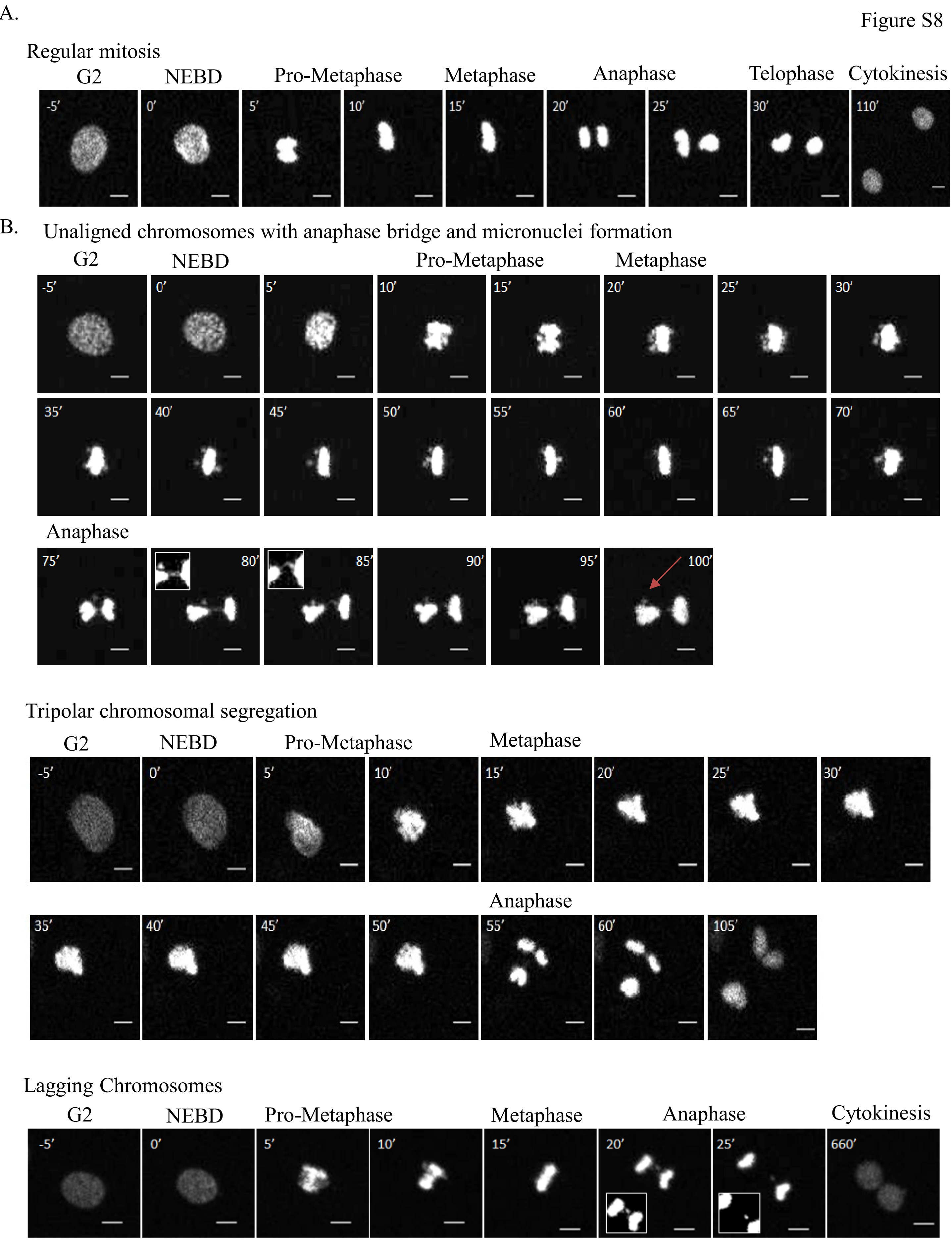

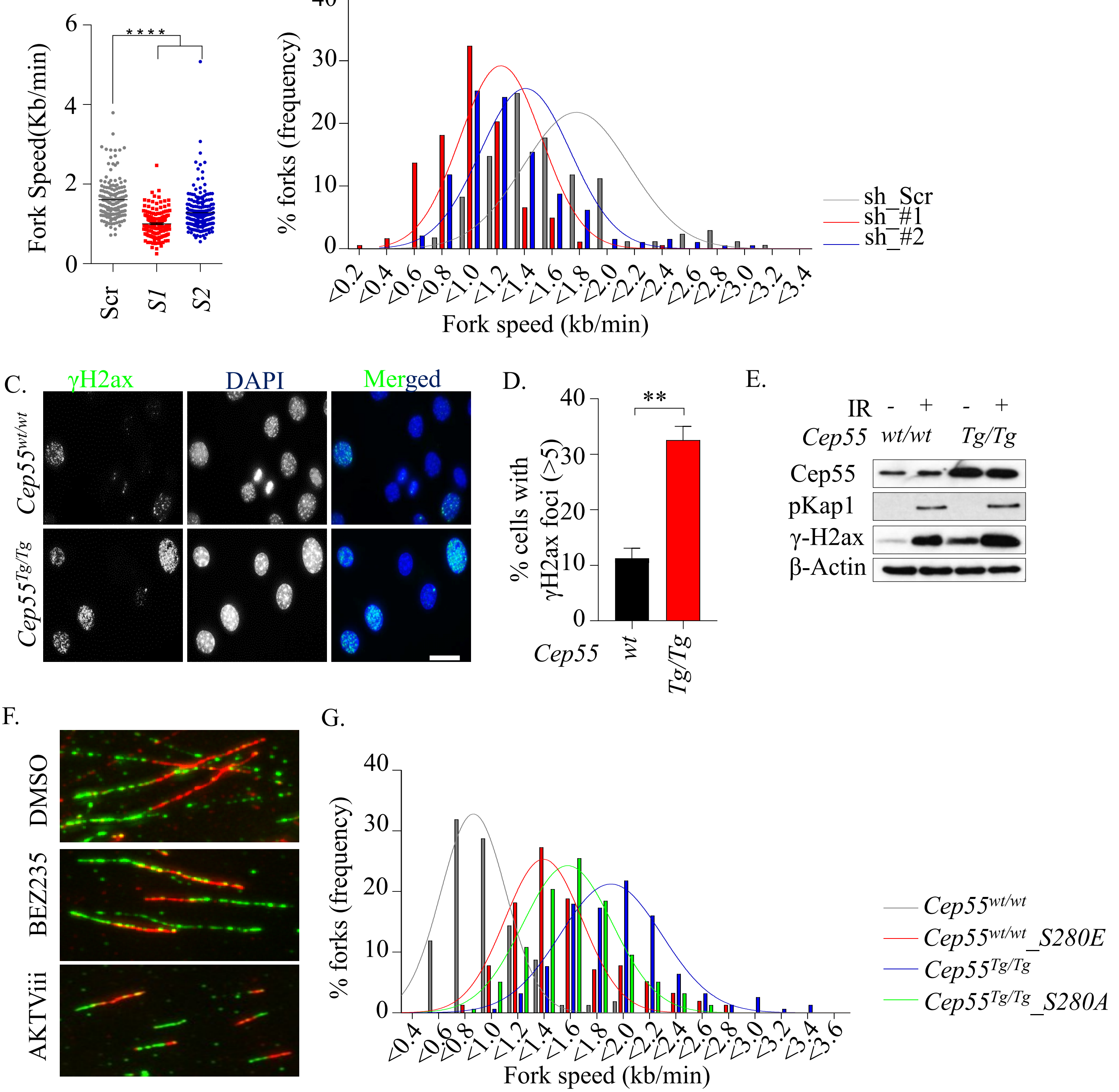
